# Human placental organoids as a model to probe early gestation maternal immune dynamics

**DOI:** 10.1101/2024.06.05.597501

**Authors:** Emily M. Slaby, Elise M. Brown, Matthew W. Becker, Nathaniel Hansen, Ritin Sharma, Patrick Pirrotte, Jessica D. Weaver

## Abstract

During pregnancy, the human placenta establishes tolerance toward fetal allogeneic tissue, where specialized trophoblast subtypes play a complex role in local and peripheral immunomodulation. However, due to inadequate models to study the early gestation of the human placenta, each trophoblast subtype’s role in modulating the maternal immune response has remained elusive. Here, we derived human placental organoids from early gestation trophoblast stem cells to (1) identify patterns of immunomodulatory protein expression by trophoblast subtype and (2) evaluate the effects of the placental organoid secretome on immune cell activation and regulation. We show that the three primary trophoblast phenotypes had distinct influences on immune cell phenotype and activation and that three-dimensional culture significantly alters trophoblast immunomodulation relative to traditional two-dimensional trophoblast culture.

## Introduction

During pregnancy, the human placenta maintains a unique, complex immune environment to support the semi- or fully allogeneic developing fetus. The placenta must balance establishing fetal antigen-specific immune tolerance with inflammatory processes, such as remodeling of the maternal decidual tissue and vasculature to support fetal nutrient needs (*1*, *2*). Specialized and immunomodulatory placental trophoblasts drive the placenta’s integration into the decidua and maintenance of tolerance at the maternal-fetal interface (*3*). Trophoblasts arise from the outside layer of the blastocyst and have primarily three distinct phenotypes, where cytotrophoblasts (CT) are the proliferative progenitor to the syncytiotrophoblast (ST) and extravillous trophoblast (EVT) phenotypes (*4*). CTs fuse to create multinucleated and syncytialized STs that provide nutrient and gas exchange between the developing embryo and maternal blood (*5*). CTs also differentiate into EVTs, which migrate into the maternal decidua to remodel the environment and promote angiogenesis (*6*). As trophoblasts act as the primary interface between the maternal decidua and developing fetus, they are critical to the development and maintenance of antigen-specific tolerance toward the fetus (*7*); however, due to inadequate models (*8*), our understanding of early gestation primary human trophoblast-immune cell interactions is limited.

The majority of studies investigating the immunomodulatory behavior of the human placenta depend on (1) animal models with physiology that exhibits limited homology to humans, (2) two-dimensional (2D) culture of late-gestation primary trophoblast cells that function differently than trophoblasts of an early gestational age and do not survive well in vitro, or (3) choriocarcinoma cell lines that do not faithfully represent healthy trophoblasts (*9*, *10*). Trophoblasts are known to express immunomodulatory surface proteins, such as human leukocyte antigen G (HLA-G), and immunomodulatory soluble factors, such as human chorionic gonadotropin subunits beta 3 and alpha (CGB3 and CGA, respectively), progesterone (P4), estrogen, placental lactogen, and relaxin (*5*, *11–13*). Though the expression of these factors has been studied at various stages of pregnancy through placenta explants and maternal blood, the complex interactions of the maternal immune system and individual trophoblast subtypes, especially at early gestation, remain enigmatic, with alternating features of inflammation and suppression throughout gestation (*14*).

Trophoblast stem cell (TSC) and TSC organoid (TO) models have recently been developed to advance the understanding of placental development and drug and virus transport across the human maternal-fetal interface (*15–20*); however, none of these models have been employed to probe human placental interactions with maternal immune cells. We previously developed a synthetic three-dimensional (3D) poly(ethylene glycol)-maleimide (PEG-mal) hydrogel platform to generate TOs comprising the three trophoblast subtypes, resulting in models corresponding to CT-, ST-, and EVT-dominated organoids (*21*). Here, we investigate how the TO secretome modulates immune cells in the maternal periphery and how culture (3D versus 2D) and differentiation (CT, ST, or EVT) conditions can alter trophoblast immunomodulatory behavior. We hypothesized that trophoblast phenotype would drive the most significant differences in immunomodulation given their distinct phenotypic protein expression and that 2D or 3D culture conditions would drive further changes in trophoblast behavior and influence over immune cells. We found that, of the three trophoblast phenotypes, the CTs induced the greatest downregulation of inflammatory cytokines and immune cell activation, and 2D and 3D culture conditions significantly modulated trophoblast influence on immune cell activation. Overall, we demonstrate that this human TO model enables the study of immune cell modulation by the trophoblast secretome, with outcomes consistent with known in vivo features of peripheral immune modulation during pregnancy.

## Results

### Trophoblast cytokine secretion is altered by 2D versus 3D culture condition

We recently developed a TO model using ECM-mimicking synthetic hydrogels as an alternative to gold-standard Matrigel culture, as the variable composition of naturally derived ECM matrices, particularly growth factors, may lead to poor interexperimental reproducibility (*22*). We demonstrated that ECM-mimicking synthetic hydrogel-generated TOs have comparable function and behavior to Matrigel-grown TOs, with synthetic PEG hydrogels driving a shift toward the ST phenotype and Matrigel driving a shift toward an EVT-predominant phenotype (*21*). Here, we sought to use this TO platform to investigate the immunomodulatory behavior of TOs polarized to a CT, ST, or EVT phenotype and compare 3D PEG- or Matrigel-grown TOs against traditional 2D culture. We hypothesized that 3D TOs would exhibit altered immunomodulatory behavior relative to 2D culture in a phenotype-dependent manner.

In our synthetic hydrogel platform, TSCs were encapsulated in PEG functionalized with collagen-derived GFOGER adhesion ligands and crosslinked with non-protease degradable (DTT) or protease degradable (VPM) linkers and cultured for 6 days with CT, ST, or EVT differentiation media (Fig. 1A). CT organoids have high expression of proliferation marker Ki67 in synthetic and Matrigel hydrogel conditions, while ST and EVT polarized organoids lose this expression after differentiation. Pan-trophoblast marker KRT7 remains highly expressed in all groups (Fig. 1B).

**Fig. 1.**
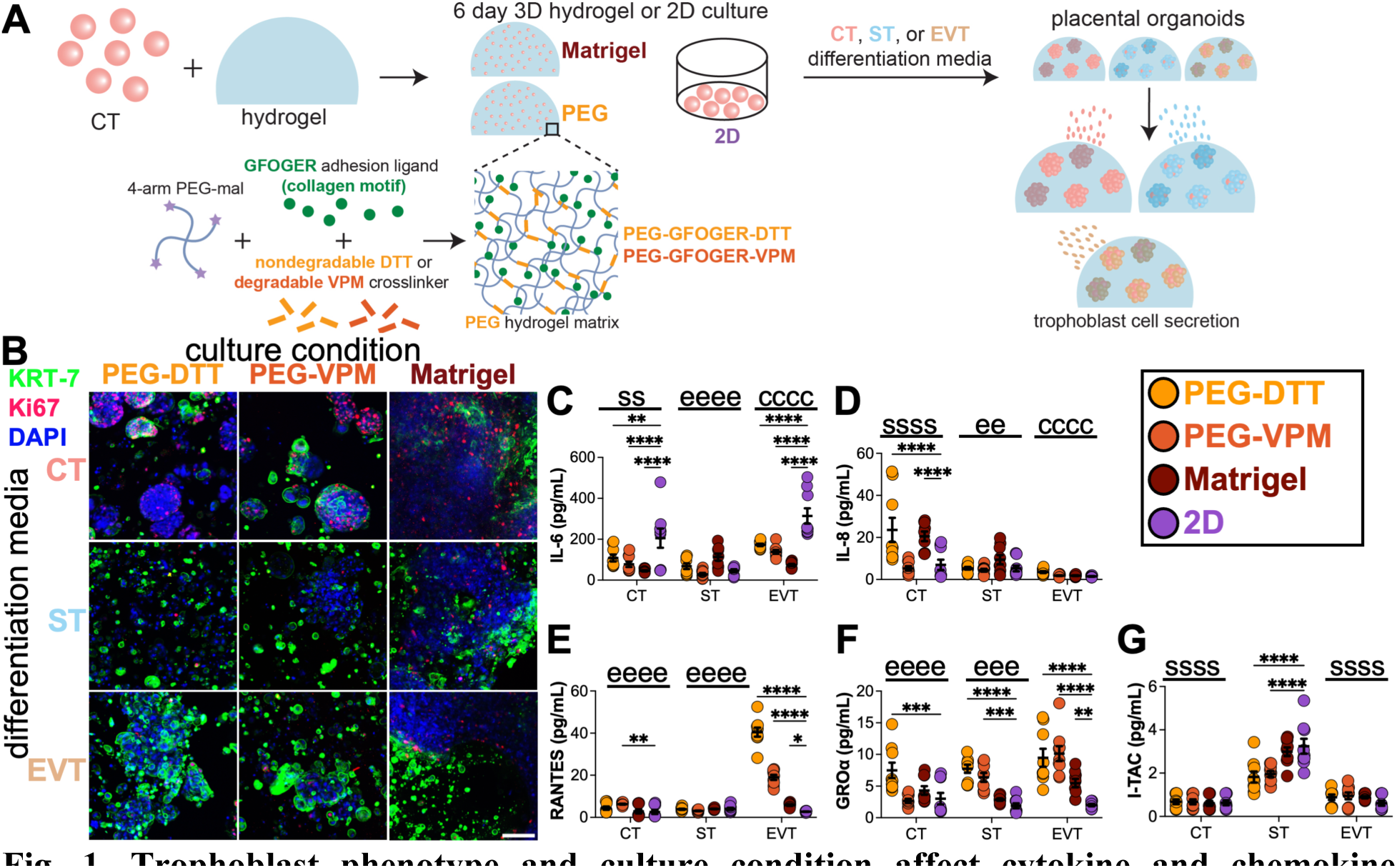
Trophoblast phenotype and culture condition affect cytokine and chemokine signaling. (**A**) TSC 1049 CT encapsulation into Matrigel or PEG hydrogels (using GFOGER adhesion ligand and either nondegradable DTT or degradable VPM crosslinker) or 2D culture for 6 days with CT, ST, or EVT differentiation media. On day 6, cells were imaged by immunocytochemistry, and supernatants were collected to analyze cytokine and chemokine secretion using LEGENDplex^TM^. (**B**) Immunocytochemistry of pan-trophoblast marker KRT7, proliferation marker Ki67, and nuclear stain DAPI and (C-G) secretion of (**C**) IL-6, (**D**) IL-8, (**E**) RANTES, (**F**) GROα, and (**G**) I-TAC from CT, ST, and EVT organoids on day 6 after culture in PEG-GFOGER-DTT, PEG-GFOGER-VPM, Matrigel, or 2D TSCs. Scale bar = 100 µm. Data are shown as mean ± SEM and analyzed by two-way ANOVA with Dunnett’s multiple comparisons test to 2D (*) and Tukey’s multiple comparisons test (c, s, and e significant to CT, ST, EVT, respectively). * p < 0.05, ** p < 0.01, *** p < 0.001, **** p < 0.0001. n=9 from 3 independent experiments.

To investigate the influence of culture condition and trophoblast phenotype on trophoblast signaling, we performed a cytokine and chemokine secretion screen on TOs or 2D-cultured TSCs (Fig. 1C-G, S1). We detected the cytokine interleukin 6 (IL-6) and chemokines IL-8 (CXCL8), RANTES (CCL5), GRO⍺ (CXCL1), and I-TAC (CXCL11). Other cytokines and chemokines were assessed but not detected (see Materials & Methods). IL-6 secretion was overall higher in 2D cultured CT and EVT differentiated cells compared to 3D (Fig. 1C). Conversely, IL-8 was highest in CT cultured in 3D PEG-DTT and Matrigel relative to 2D (Fig. 1D). RANTES and GROα secretion was significantly higher in 3D PEG conditions compared to 2D in EVT differentiated cells (Fig. 1E-F). This trend was also observed in CT and ST differentiation conditions for GROα (Fig. 1F). Generally, IL-6, RANTES, and GROα secretion was significantly higher in EVT differentiation conditions compared to CT and ST, particularly in 3D PEG culture (Fig. S1F-G, I). I-TAC secretion was significantly higher from ST differentiated cells compared to CT and EVT, with 2D culture promoting significantly higher secretion compared to PEG culture groups (Fig. 1G, S1J). Overall, 2D versus 3D culture conditions promoted significant changes in detectable cytokine and chemokine secretion.

### Trophoblast phenotype is the primary driver of differences in immunomodulatory protein expression

Trophoblasts express numerous immunomodulatory factors beyond cytokines and chemokines, many unique to the placenta. As such, we used mass spectrometry-based proteomics to comprehensively evaluate trophoblast immunomodulatory protein expression (Fig. 2, 3, S2, Table S1). CT, ST, and EVT organoids cultured in 3D hydrogels (PEG-GFOGER-VPM or Matrigel) for 6 days were compared against 2D-cultured TSC controls (Fig. 2A). Nondegradable (PEG-GFOGER-DTT) hydrogels were not included in this analysis due to inconsistent protein extraction from the hydrogel matrix. We extracted 111 proteins from our curated proteomic dataset related to the essential immunome (*23*), identifying proteins critical for innate immunity, adaptive immunity, inflammation, and the complement system (Fig. 2B, Table S1). Hierarchical clustering and principal component analysis (PCA) of these proteins demonstrated more differentially abundant protein expression across differentiation media conditions than culture condition groups (Fig. 2B-C). PCA plots illustrated that culture condition groups only clustered distinctly within the ST differentiation group, indicating the 2D versus 3D culture condition most influenced the immunome of TSCs grown under ST conditions (Fig. 2C-D). However, 3D culture groups generally clustered distinctly from 2D groups within phenotypes (Fig. 2D, bottom). Further, CT and ST Matrigel groups demonstrated unique upregulation of immune proteins characteristic of the EVT group (Fig. 2B, boxed), and the ST Matrigel group clustered closer to the EVT cluster in PCA (Fig. 2D, bottom). Overall, proteomic analysis revealed that culture and differentiation conditions significantly affect the pattern of essential immunome proteins expressed and secreted by TSCs.

**Fig. 2.**
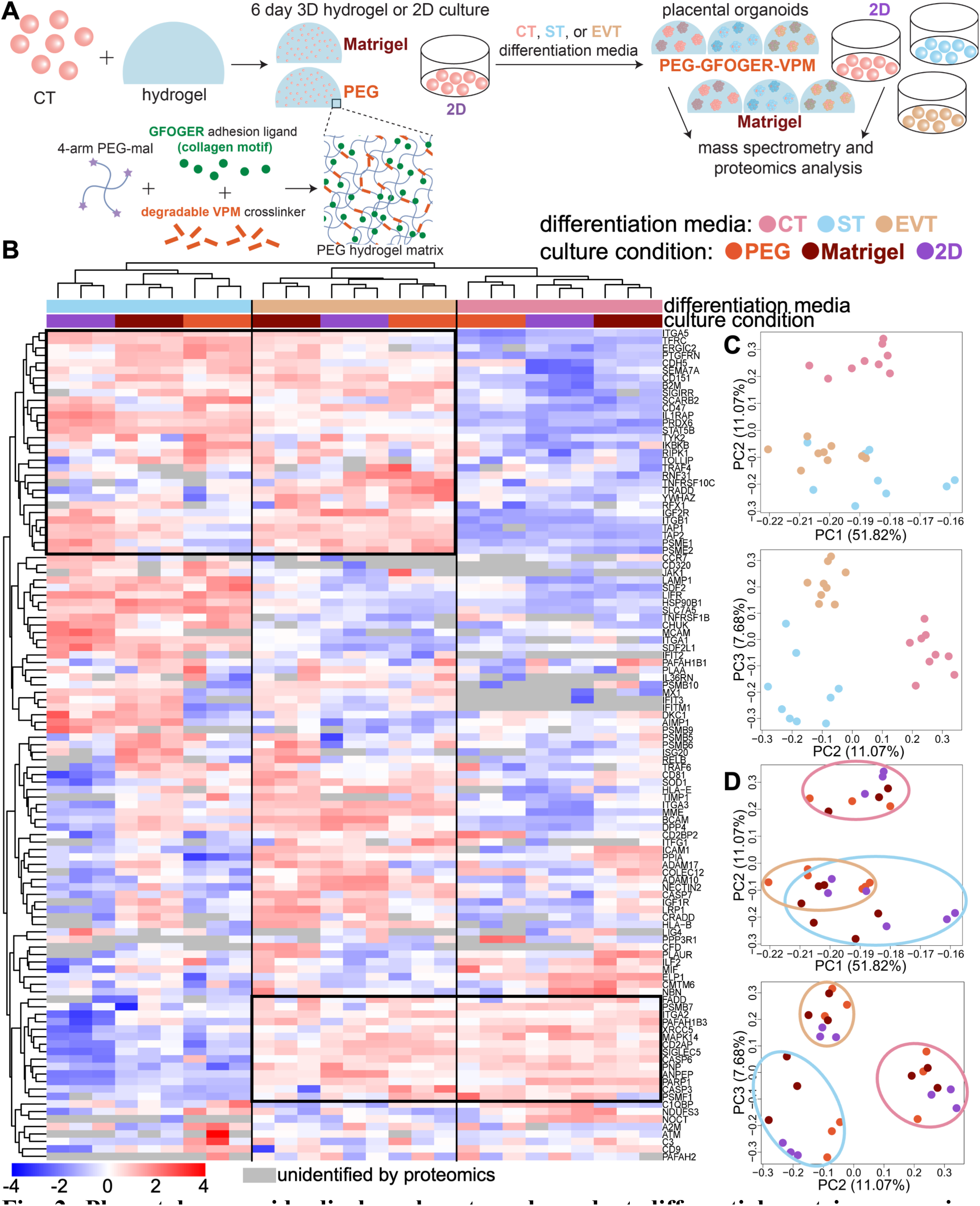
Placental organoids display phenotype-dependent differential protein expression relating to the essential immunome. (**A**) PEG-GFOGER-VPM, Matrigel, and 2D-cultured CT were cultured for 6 days in CT, ST, or EVT differentiation media and then analyzed by mass spectrometry. (**B**) Essential immunome (*23*) proteins were extracted and (**C**-**D**) clustered using principal component analysis (PCA). n=9 from 3 independent experiments.

One mechanism by which trophoblasts avoid destruction by the maternal immune system is through the surface expression of immune checkpoint molecules (Fig. 3A), which can upregulate immunosuppressive signaling in immune cells (*24*). Proteomic analysis identified T, B, natural killer (NK), and myeloid cell costimulatory molecules: programmed cell death ligand-1 (PD-L1), poliovirus receptor (PVR), CD47, poliovirus receptor-related 2 (NECTIN2, also known as PVRL2), inducible T cell costimulatory ligand (ICOSLG), and galectin 3 (LGALS3, Fig. 3B-H, S2A-H, Table S2). STs expressed the highest abundances of PD-L1 and PVR, particularly in 2D culture (Fig. 3B-C, S2A-B). STs and EVTs expressed high levels of CD47, with decreased expression in Matrigel culture (Fig. 3D, S2C). EVTs expressed the highest levels of NECTIN2 and galectins (LGALSL, LGALS8, LGALS3), and CTs expressed the highest abundances of ICOSLG and LGALS1 (Fig. 3E-H, S2D-H). Within trophoblast phenotypes, ICOSLG and LGALS3 abundances were highest in Matrigel-cultured conditions. PD-L1, PVR, and CD47 abundances were the highest in 2D conditions. HLA-E, -H, and -G, which also suppress immune cells (*12*, *25*), were highly expressed by ST and EVT phenotypes (Fig. 3I-J, S2Q-S). Overall, each trophoblast phenotype expressed checkpoint molecules that can suppress innate and adaptive immune cell activation, with expression levels significantly impacted by trophoblast phenotype and culture condition.

**Fig. 3.**
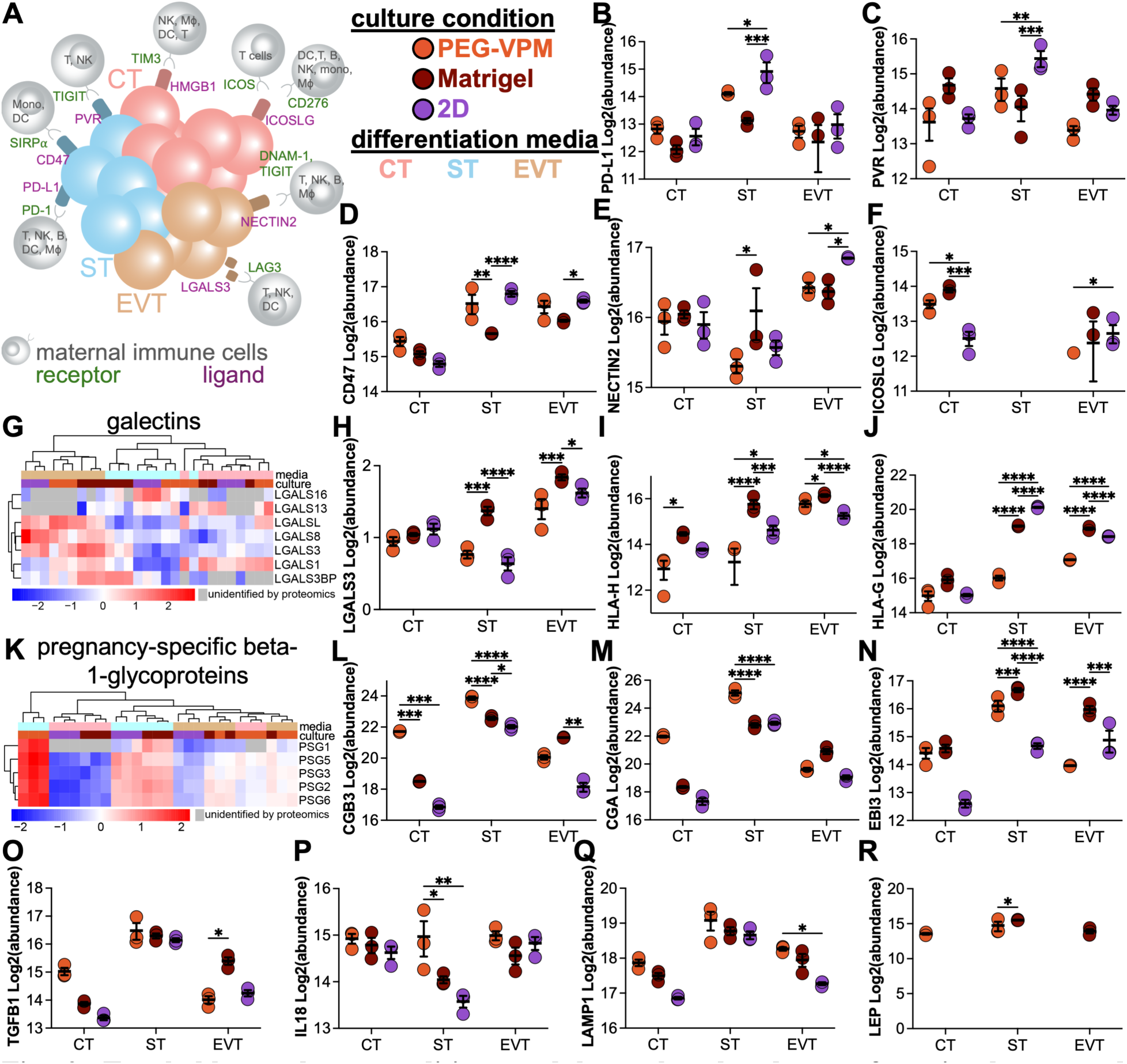
Trophoblast culture condition modulates the abundance of costimulatory and pregnancy-specific immunomodulatory proteins. (**A**) Costimulatory molecules on trophoblasts and corresponding receptors on T cells, natural killer (NK) cells, B cells, dendritic cells (DC), monocytes (mono), and macrophages (MΦ). Log2-transformed normalized abundances of expressed and secreted proteins (**B**) PD-L1, (**C**) PVR, (**D**) CD47, (**E**) NECTIN2, (**F**) ICOSLG, (**G**) galectins, (**H**) LGALS3, (**I**) HLA-H, (**J**) HLA-G, (**K**) pregnancy-specific beta-1-glycoproteins (PSG), (**L**) choriogonadotropin subunit beta 3 (CGB3), (**M**) CGA, (**N**) EBI3, (**O**) transforming growth factor beta-1 (TGFB1), (**P**) IL-18, (**Q**) lysosomal-associated membrane protein 1 (LAMP1, also known as CD107a), and (**R**) leptin (LEP). Data are shown as mean ± SEM and analyzed by two-way ANOVA with Tukey’s multiple comparisons tests. * p < 0.05, ** p < 0.01, *** p < 0.001, **** p < 0.0001. n=3.

We also identified pregnancy-related immunomodulatory molecules in our proteomics dataset. Pregnancy-specific beta-1-glycoproteins (PSG) play a role in maternal immune regulation by inducing regulatory immune cells that produce anti-inflammatory cytokines (*26–28*). Proteomics results revealed PSG1-6 and 11, CGB3 and CGA, Epstein-Barr virus-induced gene 3 (EBI3), transforming growth factor beta 1 (TGFB1), interleukin-18 (IL-18), lysosomal-associated membrane protein 1(LAMP1), and leptin (LEP) abundances were the highest in ST differentiated cells, particularly in the PEG condition, with the exceptions of EBI3 and LEP which were higher in Matrigel-cultured conditions, and TGFB1 and LAMP1, which was not significant between ST-culture conditions (Fig. 3K-R, S2I-P, Table S2). Similar to surface marker expression, secretion of pregnancy-associated immunomodulatory molecules varied by culture condition and trophoblast phenotype. Overall, we observed significant upregulation of PSGs in the ST phenotype, particularly the PEG condition, which we previously found most effectively skewed TOs toward the ST phenotype (*21*), and significant upregulation of galectins in the EVT phenotype.

### CT secretome suppresses, while ST and EVT secretome upregulates, peripheral immune cell cytokine production

After we identified distinct differences in immunomodulatory soluble factors secreted by TSCs under 2D or 3D cultured TOs and CT, ST, or EVT differentiation conditions, we were interested in the functional influence of the TSC secretome (trophoblast conditioned media, TCM) on peripheral immune cells (Fig. 4, S3). Female human peripheral blood mononuclear cells (PBMC) were exposed to TCM from each TSC group for 48 hours (Fig. 4A), without (Fig. 4B-H, S3A-M) or with the addition of activation by phorbol myristate acetate (PMA) and ionomycin at the 24-hour time point (Fig. 4I-O, S3N-Z) and assessed for inflammatory cytokine secretion. Control PBMCs were cultured with transforming growth factor beta (TGFβ) and IL-10, progesterone (P4), and human chorionic gonadotropin (hCG) as positive control soluble factors known to modulate maternal immune cells (*26*, *29–36*). Overall, unactivated PBMCs cultured in CT and EVT TCM conditions had lower inflammatory cytokine secretion than ST TCM (Fig. 4B), and EVT TCM upregulated chemotactic cytokine secretion (Fig. 4D-F). Interestingly, the TGFβ + IL-10 control condition produced the highest secretion of IL-1β, IFN-α2, MCP-1, IL-12p70, and IL-18 from unactivated PBMCs compared to P4 and hCG control conditions. This pattern was comparable to the behavior of ST TCM.

**Fig. 4.**
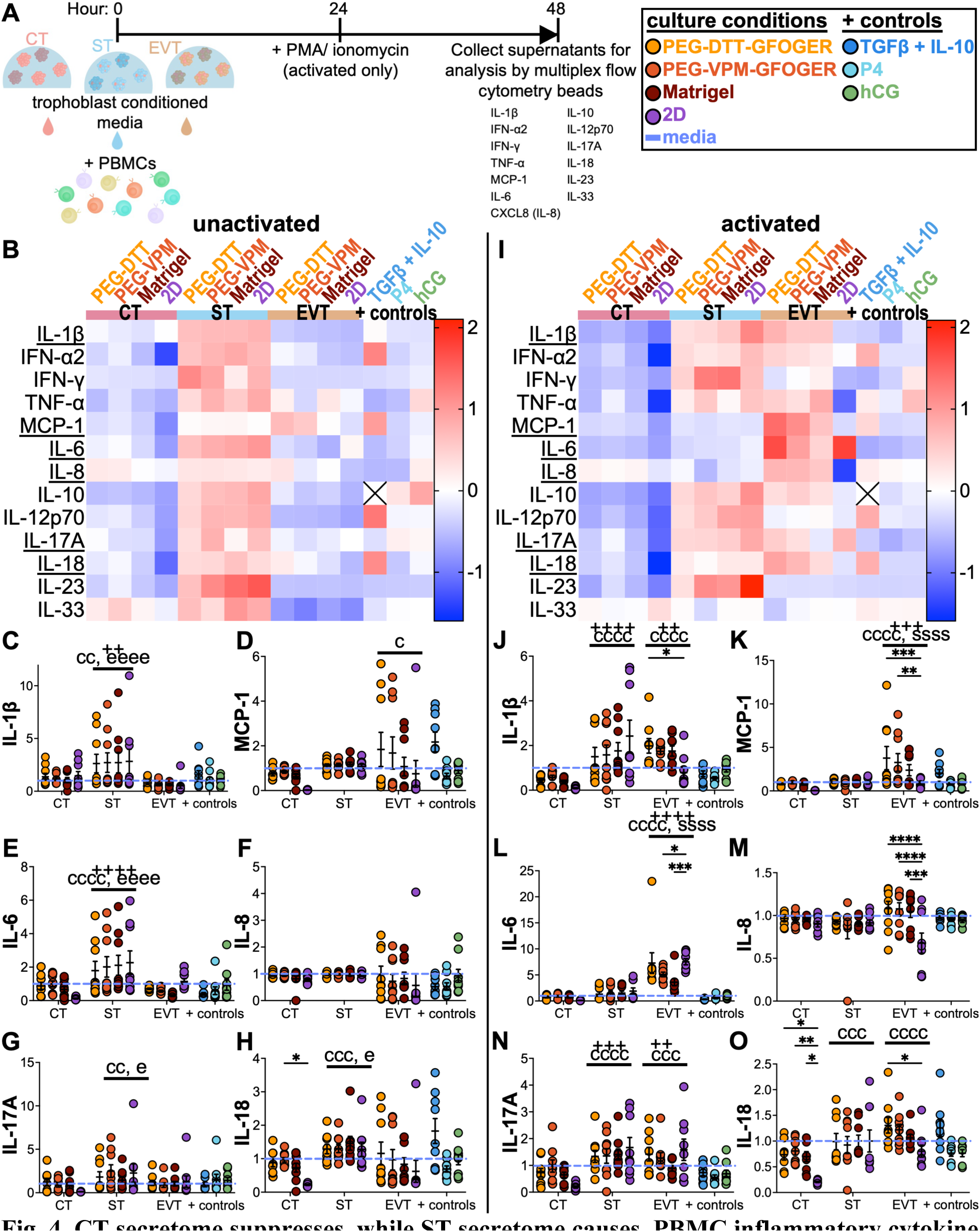
CT secretome suppresses, while ST secretome causes, PBMC inflammatory cytokine secretion in unactivated and activated conditions. (**A**) Schematic of the experimental plan: CT, ST, and EVT trophoblast conditioned media (TCM) were cultured with PBMCs for 48 hours unactivated or activated with PMA and ionomycin in the last 24 hours before collecting supernatants for analysis via LEGENDplex. (**B**) Heatmap of inflammatory cytokine secretion normalized to cell-specific media controls and standardized after a 48-hour culture with TCM or positive (+) controls. Cytokine secretion normalized to cell-specific media secretion of (**C**) IL-1β, (**D**) MCP-1, (**E**) IL-6, (**F**) IL-8, (**G**) IL-17A, (**H**) IL-18. (**I**) Heatmap of inflammatory cytokine secretion normalized to cell-specific media controls and standardized after a 48-hour culture with TCM or positive (+) controls, activated at 24 hours with positive (+) controls, and evaluated by LEGENDplex. Cytokine secretion normalized to cell-specific media secretion of (**J**) IL-1β, (**K**) MCP-1, (**L**) IL-6, (**M**) IL-8, (**N**) IL-17A, (**O**) IL-18. Data are shown as mean ± SEM and analyzed by two-way ANOVA with Dunnett’s multiple comparisons test to the 2D control (*) or Tukey’s multiple comparisons test between CT, ST, EVT, and + controls (c, s, e, +, respectively). * p < 0.05, ** p < 0.01, *** p < 0.001, **** p < 0.0001. n=9 from 3 independent experiments.

Similar to unactivated PBMCs, activated PBMCs exposed to ST and EVT TCM conditions exhibited higher inflammatory cytokine secretion than CT TCM and controls (Fig. 4I-O, S3N-Z). EVT TCM-exposed activated PBMCs also produced significantly higher secretion of MCP-1 and IL-6 than CT and ST TCM (Fig. 4K-L, S3R-S). Interestingly, PBMC activation only significantly altered cytokine secretion trends in the EVT TCM-exposed condition, primarily in 3D TO conditions, except IL-6 and IL-17, where 2D caused the highest secretion (Fig. 4l, 4N, S3S, S3V). Overall, PBMC cytokine secretion was mainly impacted by trophoblast phenotype, with each phenotype producing a unique expression pattern and 2D versus 3D culture condition exhibiting the greatest influence in CT and EVT phenotypes. CT-exposed PBMCs exhibited relatively reduced cytokine secretion, with the greatest reduction in the 2D culture group, and EVT-exposed PBMCs exhibited increased cytokine secretion in primarily the 3D cultured trophoblast conditions.

### Trophoblast secretome alters the activation profile of human NK and T cell lines in a phenotype- and culture condition-dependent manner

With the observation that TSC phenotypic differentiation and culture condition can influence PBMC cytokine secretion, we next aimed to determine the effects that the TSC secretome could have on the phenotype and activation of immune cells in the maternal periphery. We first used the NK and T cell lines, NK-92 and Jurkat, respectively, to determine if trophoblast secretome from varying differentiation media and culture type (2D versus 3D) would affect NK and T cell phenotype. NK-92 and Jurkats were cultured with TCM (26 or 48 hr, respectively) from CT, ST, or EVT differentiated TSCs cultured in 3D (PEG-GFOGER-DTT, PEG-GFOGER-VPM, and Matrigel) or 2D for 6 days and phenotypic changes were evaluated by flow cytometry (Fig. S4A, S5A). Positive controls were pregnancy-specific immunomodulators TGFβ, TGFβ and IL-10, P4, and hCG. While TCM from CT did not significantly affect the percentage of LAMP-1^+^ (CD107a, degranulation marker) unactivated or activated NK-92s, CT TCM reduced IFNγ secretion from unactivated NK-92s, and 2D CT TCM reduced IFNγ secretion from activated NK-92s comparably to TGFβ controls (Fig. S4B-E). Overall, TCM reduced NK cell line secretion of IFN-γ in a culture condition-specific manner.

In unactivated and activated Jurkats, CT, ST, and EVT TCM significantly increased the expression of activation markers CD38 and CD71 (CT and ST only for unactivated) compared to media controls, whereas CT TCM significantly decreased the expression of CD154 (also known as CD40L) compared to media controls and ST and EVT TCM (Fig. S5B-O). Unactivated Jurkats cultured with 2D TCM had significantly increased expression of CD38 (EVT), CD69 (CT, EVT), and CD71 (CT) compared to 3D TCM (Fig. S5B-H). The same trend was seen in activated Jurkats with the expression of CD38 (CT, EVT) and CD71 (CT, Fig. S5I-O). Conversely, activated Jurkats cultured with 2D TCM had significantly decreased expression of CD25 (CT), CD69 (CT), and CD154 (CT, EVT). Overall, TCM altered Jurkat T cell line phenotypic activation markers in a culture condition- and trophoblast phenotype-dependent manner, with the greatest reductions observed for the CT phenotype.

We saw apparent differences in NK and T cell line activation in response to TCM generated under different culture and differentiation conditions; however, these observations were in isolated immune cell types, which does not account for interactions between immune cells as would occur natively in vivo. As such, we next evaluated how individual immune cell phenotypes are altered in response to TCM in a mixed leukocyte environment, using primary human female PBMCs. We observed phenotypic shifts after TCM exposure (denoted as unactivated) and after PBMC activation with PMA and ionomycin to simulate TCM impact on activated peripheral maternal immune cells. We evaluated phenotypic changes in key immune cells in pregnancy: NK cells (cytotoxic CD16^+^ CD56^dim^ or cytokine-secreting CD16^-^ CD56^++(bright)^), T cells (T helper CD3^+^ CD8^-^ or cytotoxic CD3^+^ CD8^+^), and B cells (CD19^+^). All expression levels (MFI) are normalized to CT, ST, or EVT media controls with standardization to show upregulation or downregulation of each phenotypic marker. Time points were chosen according to preliminary studies that identified the highest activation marker levels after PMA and ionomycin addition.

### Trophoblast secretome produces contrasting activation profiles in cytotoxic and cytokine-secreting NK cell phenotypes in a trophoblast phenotype-dependent manner

CD16^+^ CD56^dim^ and CD16^-^ CD56^++^ NK cells are two major NK subsets in the peripheral blood and are considered more cytotoxic or cytokine-secreting, respectively (*37*, *38*). In unactivated PBMCs, greater than 85% of cytotoxic CD16^+^ CD56^dim^ NK cells (Fig. 5A, S6-8), which typically make up 90% of peripheral NKs (*39*, *40*), expressed Granzyme B (Fig. S7A-B). However, CT TCM resulted in a significantly higher percentage of Granzyme B-expressing cells than ST or EVT TCM (Fig. S7B), while the 2D CT TCM reduced Granzyme B MFI relative to 3D culture (Fig. 5B-C, S8A). Additionally, positive controls and ST TCM produced the highest expression of markers of cytotoxic activity, Granzyme B and CD107a (Fig. 5B-D, S8A-B), respectively. Similarly, CD158a and CD38, which are also involved with cytotoxic activity, exhibited the highest expression on unactivated cytotoxic CD16^+^ CD56^dim^ NK cells cultured in EVT TCM and 2D CT TCM (Fig. 5B, S8C-D). Conversely, activated cytotoxic CD16^+^ CD56^dim^ NK cells exhibited more consistent expression of cytotoxic activity markers within trophoblast phenotype groups, where Granzyme B, CD107a, CD158a, and CD38 were upregulated in CT and EVT TCM culture conditions, with ST TCM-exposed cytotoxic NKs expressing significantly lower CD107a and CD38 than CT and EVT TCM conditions (Fig. 5E-G, S8E-H). Few significant differences were observed in the percentage of cells positive for these markers (Fig. S7F-J).

**Fig. 5.**
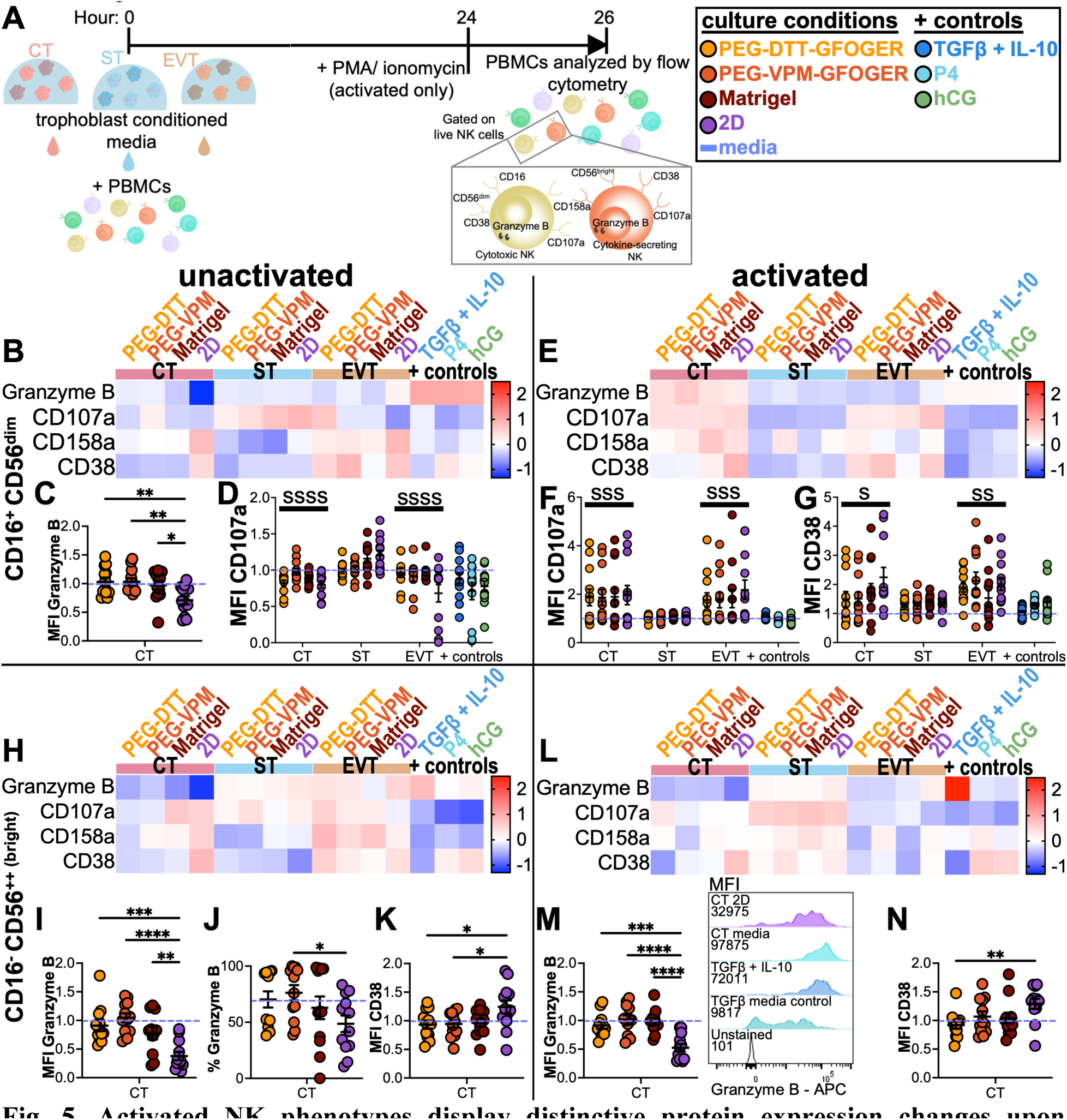
Activated NK phenotypes display distinctive protein expression changes upon exposure to trophoblast secretome. (**A**) Experimental setup: PBMCs were cultured with TCM for 26 hours with or without the addition of activation with PMA and ionomycin after 24 hours, then analyzed by flow cytometry according to the gating strategy in Fig. S6. (**B**) Heatmap of unactivated cytotoxic NK cell (CD16^+^ CD56^dim^) protein expression normalized to cell-specific media controls and standardized, (**C**) Granzyme B MFI, and (**D**) CD107a MFI after a 26-hour culture of PBMC with TCM or controls. (**E**) Heatmap of activated cytotoxic NK cells (CD16^+^ CD56^-^) protein expression normalized to cell-specific media controls and standardized after a 26-hour culture with TCM or controls, activated at 24 hours, (**F**) CD107a MFI, and (**G**) CD38 MFI. (**H**) Heatmap of unactivated cytokine-secreting NK cells (CD16^-^ CD56^++^) protein expression normalized to cell-specific media controls and standardized after a 26-hour culture of PBMC with TCM or controls, (**I**) Granzyme B MFI, (**J**) percentage of Granzyme B^+^ cells, and (**K**) CD38 MFI. (**L**) Heatmap of activated cytokine-secreting NK cells (CD16^-^ CD56^++^) protein expression normalized to cell-specific media controls and standardized after a 26-hour culture with TCM or controls, activated at 24 hours, (**M**) Granzyme B MFI with representative flow cytometry plots, and (**N**) CD38 MFI. Data are shown as mean ± SEM and analyzed by ordinary one-way ANOVA with Dunnett’s multiple comparisons test to the 2D control (*) or (D, F, G) ordinary two-way ANOVA with main effects only and Tukey’s multiple comparisons test (s significant to ST). * p < 0.05, ** p < 0.01, *** p < 0.001, **** p < 0.0001. n=12 from 4 independent experiments.

The cytokine-secreting CD56^++^ NK cell phenotype is typically a minority 10% in peripheral blood (*39*, *40*), is the predominant NK phenotype found in lymph nodes, and represents 70-80% of lymphocytes in the early pregnancy decidua (*41*). CD56^++^ NK cells had markedly different patterns of activation marker expression relative to the cytotoxic CD16^+^ CD56^dim^ NK phenotype (Fig. 5H-N, S6-8). Granzyme B expression was the lowest in unactivated CD56^++^ cells cultured in CT TCM, where the 2D cultured CTs significantly reduced Granzyme B MFI and percentage positive relative to 3D culture (Fig. 5H-J, S7K-L, S8I). Generally, unactivated CD56^++^ NK cells cultured in EVT TCM had the highest expression of activation markers Granzyme B, CD107a, CD158a, and CD38 (Fig. 5H, S8I-L), while 2D CT TCM significantly upregulated CD38 relative to 3D culture (Fig. 5K, S8L). Conversely, activated CD56^++^ NK cells exhibited lower expression of activation markers when cultured with EVT TCM compared to ST TCM (Fig. 5L). Similar to unactivated CD56^++^ NK, activated CD56^++^ NK cells in 2D CT TCM had lower Granzyme B expression relative to 3D conditions (Fig. 5M, S8M) but higher expression of CD38 relative to PEG-DTT CT TCM (Fig. 5N, S8P). For both activated and unactivated cytokine-secreting CD56^++^ NK cells, CT TCM-exposed cells exhibited a higher percentage of positivity for Granzyme B relative to EVT TCM and reduced CD38 percentage relative to ST and EVT TCM (Fig. S7K-L, S7O-Q, S7T). Overall, trophoblast phenotype had the most significant impact on NK cell activation, and CT TCM exhibited the most variation by 2D versus 3D culture condition. Further, unactivated CD16^+^ CD56^dim^ and CD56^++^ NKs exhibited comparable activation markers when exposed to TCM media, whereas activated CD16^+^ CD56^dim^ and CD56^++^ NK exhibited contrasting activation profiles.

### ST and EVT secretome upregulates, while the CT secretome downregulates, cytotoxic and helper T cell activation and regulatory markers

We next evaluated how 2D or 3D cultured TCM influenced T helper cell activation within our PBMC population (Fig. 6, S9-11). Upon activation with PMA and ionomycin, CD4 is rapidly downregulated from the T cell surface and endocytosed (*42*); therefore, for consistency of gating strategy between unactivated and activated groups, CD8^+^ and CD8^-^ populations are labeled as cytotoxic and T helper cells, respectively (Fig. S9). Unactivated T helper cells did not exhibit a clear pattern of trophoblast phenotype-dependent activation in response to TCM, except for FOXP3, which was downregulated with CT TCM, relatively unchanged in ST TCM, and upregulated with EVT TCM (Fig. 6B, S11E). There were significant differences in 2D versus 3D culture conditions within trophoblast phenotypes, where 2D CT TCM resulted in a significant increase in the percentage of CD69^+^ T helper cells (Fig. 6C, S10G) and the MFI of CD69 (Fig. 6D, S11F) and CD38 (Fig. 6E, S11D) and a significant decrease in the MFI of CD154 (Fig. 6F, S11C), Granzyme B (Fig. 6G, S11A), and FOXP3 (Fig. 6H, S11E). 2D EVT TCM also significantly reduced FOXP3 MFI relative to 3D (Fig. 6H, S11E).

**Fig. 6.**
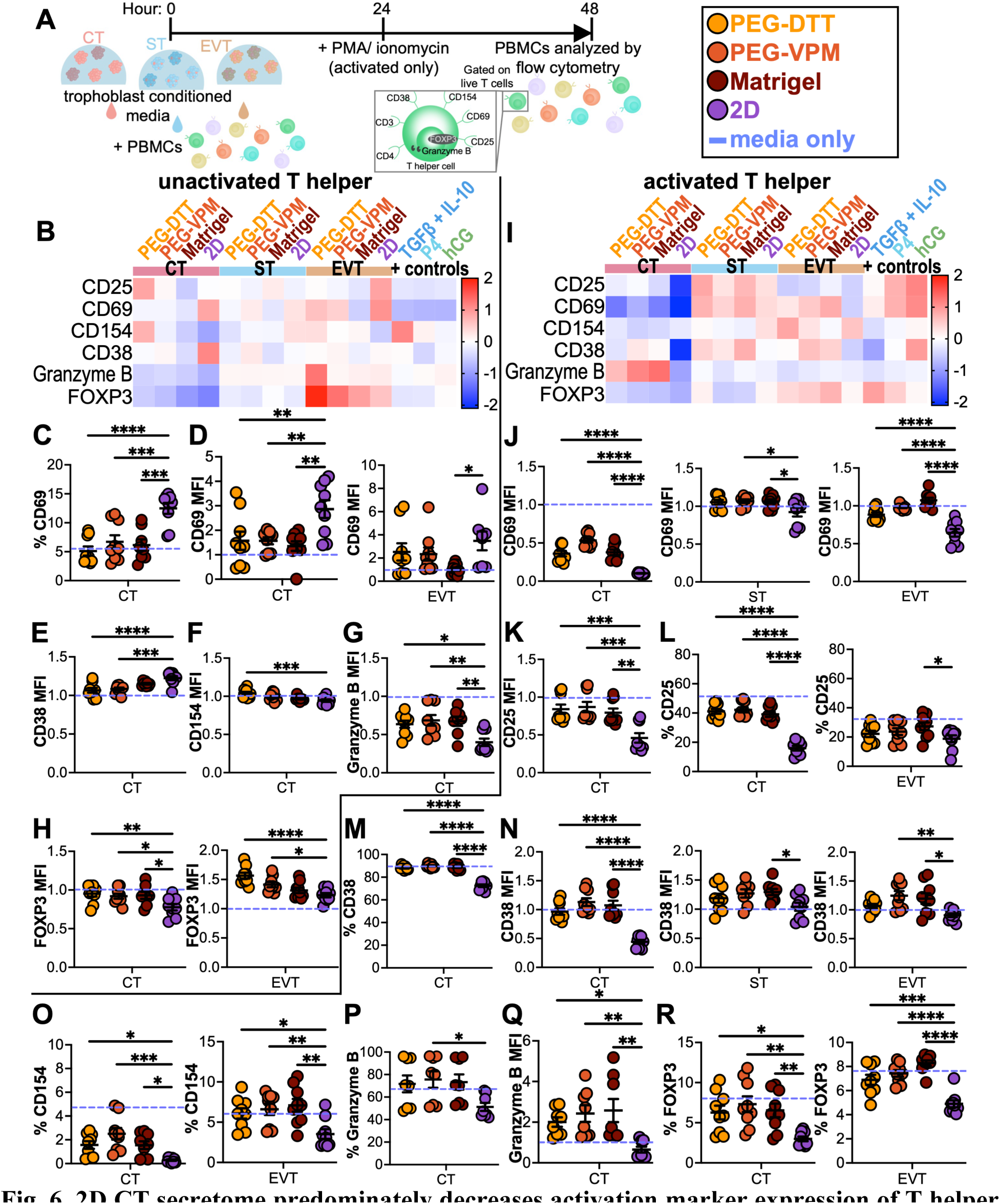
2D CT secretome predominately decreases activation marker expression of T helper cells within PBMCs compared to 3D CT secretome. (**A) Experimental layout: PBMCs were** cultured with TCM for 48 hours with or without activation by PMA and ionomycin in the last 24 hours before analysis of the T helper cell (CD3^+^ CD8^-^) population using flow cytometry (gating strategy in Fig. S9). (**B**) Heatmap of unactivated T helper cell protein expression normalized to cell-specific media controls and standardized after a 48-hour culture of PBMC with TCM or controls, (**C**) percentage of CD69^+^ cells, (**D**) CD69 MFI, (**E**) CD38 MFI, (**F**) CD154 MFI, (**G**) Granzyme B MFI, and (**H**) FOXP3 MFI. (**I**) Heatmap of activated T helper cell protein expression normalized to cell-specific media controls and standardized after a 48-hour culture with TCM or controls, activated with PMA and ionomycin at 24 hours, (**J**) CD69 MFI, (**K**) CD25 MFI, (**L**) percentage of CD25^+^ cells, (**M**) percentage of CD38^+^ cells, (**N**) CD38 MFI, (**O**) percentage of CD154^+^ cells, (**P**) percentage of Granzyme B^+^ cells, (**Q**) Granzyme B MFI, and (**R**) percentage of FOXP3^+^ cells after incubation. Data are shown as mean ± SEM and analyzed by ordinary one-way ANOVA with Dunnett’s multiple comparisons test to the 2D control. * p < 0.05, ** p < 0.01, *** p < 0.001, **** p < 0.0001. n=9 from 3 independent experiments.

Activated T helper cells exhibited more apparent differences in activation markers between trophoblast phenotypes, where CT TCM decreased CD69 and increased Granzyme B relative to ST and EVT TCM (Fig. 6I). Notably, ST and EVT activation markers were fairly aligned with P4 and hCG controls. In contrast, the CT phenotype exhibited a contrasting activation to these controls. CD69 expression strongly depended on trophoblast 2D or 3D culture conditions, where 2D culture reduced CD69 MFI relative to 3D in CT, ST, and EVT TCM (Fig. 6J, S11L). Similarly, 2D TCM caused a reduction in CD25 MFI with CT TCM (Fig. 6K, S11H) and the percentage of CD25^+^ cells with 2D CT and EVT TCM (Fig. 6L, S10J) compared to 3D TCM. Additionally, CD38, typically upregulated in T cells during pregnancy (*43*), exhibited a reduced percentage of CD38^+^ cells with 2D CT CTM (Fig. 6M, S10L) and CD38 MFI with 2D TCM relative to 3D conditions (Fig. 6N, S11J). This pattern of reduced activation in 2D cultured cells was also observed in the percentage of CD154^+^ T helper cells with culture in CT and EVT TCM (Fig. 6O, S10K), the percentage of Granzyme B^+^ T helper cells with CT TCM (Fig. 6P, S10I), Granzyme B MFI (Fig. 6Q, S11G) with CT TCM, and the percentage of FOXP3^+^ cells, indicative of a regulatory T cell (Treg) phenotype, with CT and EVT TCM (Fig. 6R, S10M). Generally, the percentage of cells positive for activation markers in activated T helper cells was significantly higher when cultured in CT TCM compared to ST and EVT TCM (Fig. S10H-N). Further, PBMC activation led to an increase in the percentage of CD25^+^ FOXP3^+^ Tregs in all groups, including media controls, apart from 2D CT and EVT TCM, where this increase was suppressed (Fig. S10O). Overall, TCM modulation of T helper cell activation had a high dependence on trophoblast 2D or 3D culture conditions, and the CT phenotype resulted in a distinctive phenotype relative to ST and EVT.

We also evaluated phenotypic shifts in the cytotoxic CD8^+^ T cell population (Fig. 7, S9, S12-13). Unactivated CD8^+^ T cells exhibited moderate phenotype-dependent differences in activation, with markers such as Granzyme B broadly downregulated in with CT TCM and FOXP3 expression broadly upregulated with EVT TCM (Fig. 7B). Again, 2D TCM exhibited significant differences from 3D TCM in cytotoxic T cells, where 2D CT CD69^+^ percentage (Fig. 7C, S12G), CD69 MFI for 2D CT and EVT (Fig. 7D, S13F), and CD38 MFI for 2D CT (Fig. 7E, S13D) was significantly higher than when cultured with 3D TCM. Conversely, Granzyme B (Fig. 7F, S13A) and CD25 MFI (Fig. 7G, S13B) were lower with 2D CT TCM culture compared to 3D CT TCM. FOXP3 expression was significantly higher in CD8^+^ cells cultured with EVT TCM compared to CT and ST TCM (Fig. 7H, S13E). Within unactivated cytotoxic T cells, CT TCM significantly modulated the percentage of positive cells with these activation markers compared to ST and EVT TCM (Fig. S12A-G).

**Fig. 7.**
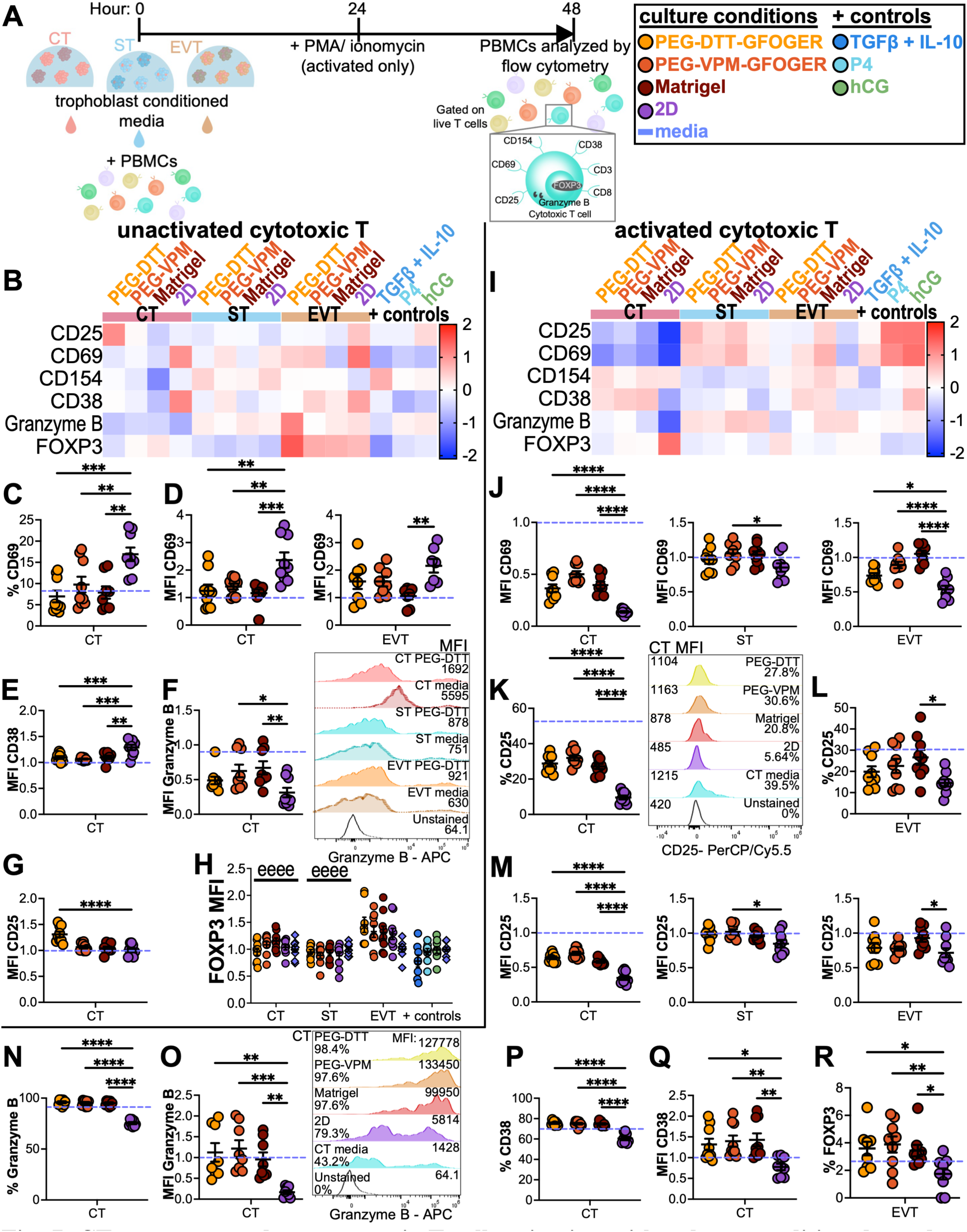
CT secretome reduces cytotoxic T cell activation with culture condition-dependent responses. (**A**) Experimental layout: PBMCs were cultured with TCM for 48 hours with or without activation by PMA and ionomycin in the last 24 hours before analysis of the cytotoxic T cell (CD3^+^ CD8^+^) via flow cytometry (gating strategy in Fig. S9). (**B**) Heatmap of unactivated cytotoxic T cell protein expression normalized to cell-specific media controls and standardized after a 48-hour culture of PBMC with TCM or controls, (**C**) percentage of CD69^+^ cells, (**D**) CD69 MFI, (**E**) CD38 MFI, (**F**) Granzyme B MFI with representative flow cytometry plots, (**G**) CD25 MFI, and (**H**) FOXP3 MFI. (**I**) Heatmap of activated cytotoxic T cell protein expression normalized to cell-specific media controls and standardized after a 48-hour culture with TCM or controls and activated with PMA and ionomycin at 24 hours, (**J**) CD69 MFI, (**K**-**L**) the percentage of CD25^+^ cells with representative flow cytometry plots, (**M**) CD25 MFI, (**N**) the percentage of Granzyme B^+^ cells, (**O**) Granzyme B MFI with representative flow cytometry plots, (**P**) the percentage of CD38^+^ cells, (**Q**) CD38 MFI, and (**R**) the percentage of FOXP3^+^ cells. Data are shown as mean ± SEM and analyzed by ordinary one-way ANOVA with Dunnett’s multiple comparisons test to the 2D control or (H) ordinary two-way ANOVA with main effects only and Tukey’s multiple comparisons test (e significant to EVT). * p < 0.05, ** p < 0.01, *** p < 0.001, **** p < 0.0001. n=9 from 3 independent experiments.

Activated cytotoxic T cells from the PBMC population exhibited a more robust pattern of trophoblast phenotype-dependent activation (Fig. 7I), where CT TCM-exposed cytotoxic T cells expressed significantly lower CD25 and CD69 expression compared to ST and EVT TCM (Fig. S13H, S13L). CD69 MFI (Fig. 7J, S13L), percentage of CD25^+^ cells (Fig. 7K-L, S12J), and CD25 MFI (Fig. 7M, S13H) tended to be lower with 2D cultured TCM relative to 3D TCM in nearly all phenotype conditions. A similar pattern of reduced activation markers in the 2D condition was observed for the percentage of Granzyme B^+^ cells (Fig. 7N, S12I) and MFI (Fig. 7O, S13G) with CT TCM, the percentage of CD38^+^ cells (Fig. 7P, S12L) and MFI (Fig. 7Q, S13J) with CT TCM, and the percentage of FOXP3^+^ T cells exposed to EVT TCM (Fig. 7R, S12M). While CT TCM exhibited the most significant modulation of activation markers, ST and EVT TCM also showed significant modulation of the percentage of T cells positive for activation markers (Fig. S12H-N). Overall, trophoblast phenotype and culture condition significantly influenced the expression of T cell activation and regulatory markers in the PBMC population, with activated cytotoxic T cells exposed to CT TCM broadly downregulating activation and regulatory markers.

### Trophoblast secretome-conditioned B cells exhibit clear trophoblast phenotype-dependent phenotypic shift

Finally, B cells are known to be a key driver of tolerance in pregnancy (*44*), so we investigated the influence of TCM on CD19^+^ B cell activation within our PBMC population (Fig. 8, S14-16). In unactivated B cells, CT, ST, and EVT TCM culture conditions resulted in distinct B cell activation marker expression (Fig. 8B). CT TCM upregulated CD86 and HLA-DR MFI, ST TCM upregulated CD71 and CD40 MFI, and EVT TCM upregulated CD24, CD80, and Granzyme B MFI (Fig. 8B, S16A-G). 2D versus 3D culture differences were evident with CT TCM in CD71 percentage (Fig. 8C, S15E), HLA-DR MFI (Fig. 8D, S16B), and CD40 MFI (Fig. 8E, S16G), and CD86^+^ percentage (Fig. 8F, S15D) and MFI (Fig. 8G, S16C).

**Fig. 8.**
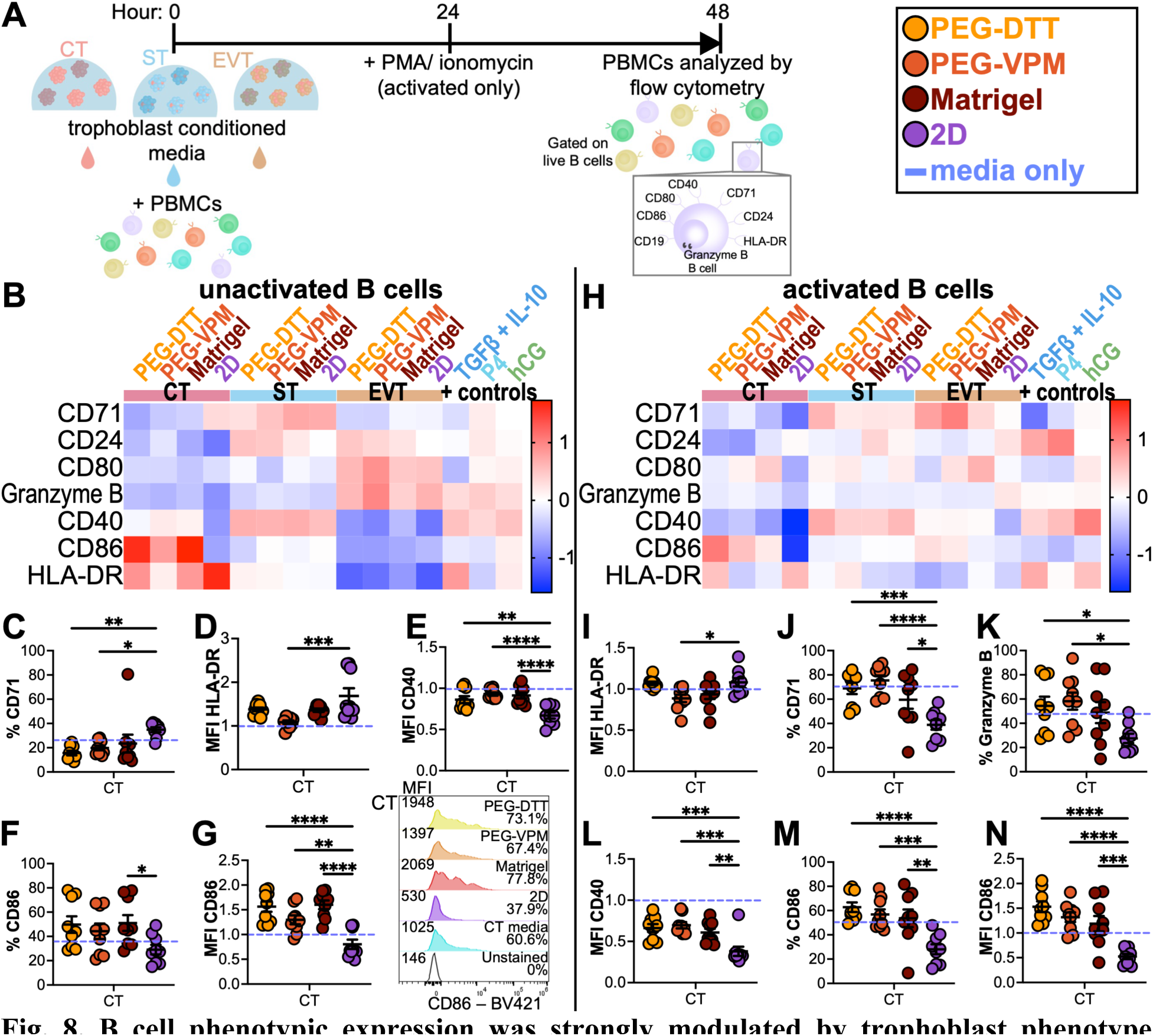
B cell phenotypic expression was strongly modulated by trophoblast phenotype secretome in unactivated cells and exhibited greater variability by 2D versus 3D culture secretome in activated B cells. (**A**) Experimental layout: PBMCs were cultured with TCM for 48 hours with or without activation by PMA and ionomycin in the last 24 hours before analysis of the B cell population (CD19^+^) via flow cytometry (gating strategy in Fig. S14). (**B**) Heatmap of unactivated B cell protein expression normalized to cell-specific media controls and standardized after a 48-hour culture of PBMC with TCM or controls, (**C**) percentage of CD71^+^ cells, (**D**) HLA-DR MFI, (**E**) CD40 MFI, (**F**) percentage of CD86^+^ cells, and (**G**) CD86 MFI with representative flow cytometry plots. (**H**) Heatmap of activated B cell protein expression normalized to cell-specific media controls and standardized after a 48-hour culture with TCM or controls, activated at 24 hours, (**I**) percent of Granzyme B^+^ cells, (**J**) percent of CD71^+^ cells, (**K**) HLA-DR MFI, (**L**) CD40 MFI, (**M**) percent of CD86^+^ cells, and (**N**) CD86 MFI of CD19^+^ cells. Data are shown as mean ± SEM and analyzed by ordinary one-way ANOVA with Dunnett’s multiple comparisons test to the 2D control. * p < 0.05, ** p < 0.01, *** p < 0.001, **** p < 0.0001. n=9 from 3 independent experiments.

Under activation conditions, B cell activation marker expression became less defined by trophoblast phenotype (Fig. 8H), though CT TCM broadly remained suppressive of CD71, CD24, and CD40 MFI. Additionally, the CT phenotype exhibited 2D versus 3D culture condition differences in HLA-DR MFI (Fig. 8I, S16I), the percentage of CD71^+^ (Fig. 8J, S15N) and Granzyme B^+^ B cells (Fig. 8K, S15K), CD40 MFI (Fig. 8L, S16N), the percentage of CD86^+^ B cells (Fig. 8M, S15M), and CD86 MFI (Fig. 8N, S16J). Generally, CT TCM significantly modulated the percentage of Granzyme B^+^ and CD86^+^ B cells in unactivated and activated conditions (Fig. S15B, S15D, S15K, S15M). Overall, B cell marker expression was strongly modulated by trophoblast phenotype in unactivated B cells and exhibited greater variability by 2D versus 3D culture condition in activated B cells.

## Discussion

Trophoblasts drive fetal tolerance by secreting immunomodulatory hormones, cytokines, chemokines, and surface presentation of immunomodulatory proteins. However, limited studies have investigated these mechanisms in a trophoblast phenotype-dependent manner. Traditionally, inadequate rodent models and 2D cultures of primary human trophoblasts or human choriocarcinoma cell lines have been used to study human placental immunological mechanisms (*9*, *10*). The recent development of human TSC and TO models may present a more faithfully representative model to study human placental immunological mechanisms (*15–20*). Organoid systems better replicate the native tissue microenvironment with 3D matrix cues and increased cell-to-cell contact. We recently demonstrated that TSC-derived placental organoids can be generated using synthetic PEG-based hydrogel comparably to the gold standard Matrigel hydrogel system (*21*), and here we employed this model to promote more physiologically relevant trophoblast behavior and evaluate trophoblast secretome impact on peripheral immune cell behavior (Fig. 1).

We profiled the CT, ST, and EVT secretome via proteomics and multiplex cytokine assays, identifying numerous immunomodulatory proteins, cytokines, and chemokines (Fig. 1-3). Among the chemokines identified, RANTES is involved in decidualization, immune regulation, and trophoblast migration at the maternal-fetal interface. Accordingly, we found the highest levels of secretions from the EVT subtype, where PEG promoted the highest levels of RANTES secretion (Fig. 1E). Patients with recurrent spontaneous abortions (RSA) and women without previous pregnancies have comparable levels of RANTES in their sera, which is significantly lower compared to women who previously had successful pregnancies. Further, RANTES can specifically induce maternal immune tolerance towards allogeneic cells in a mixed lymphocyte reaction (MLR) (*45*), which expands on studies that found pregnant serum caused proliferation suppression in an MLR (*46*). We also observed GROα in the trophoblast secretome, a chemokine to neutrophils, monocytes, T cells, and dendritic cells (DC) involved in immune regulation, angiogenesis, and inflammation. GROα was highly secreted by EVT, which also participates in angiogenesis (*47*, *48*), and had higher secretion from 3D PEG conditions in CT, ST, and EVT compared to 2D (Fig. 1F). GROα has demonstrated suppressive potential in glioblastoma and small cell lung cancer, where it is associated with the recruitment of Tregs (*49*) and could also be contributing to the increased Treg phenotypes from EVT TCM (Fig. 6B, 6I, 7B, S11E, S11K, S13E, S13K) (*50*). Proteomics results also revealed distinct phenotypic differences in pregnancy-related immunomodulatory proteins, such as PSGs, which play a role in maternal immune regulation by inducing anti-inflammatory cytokines (*26–28*). Additionally, PSGs have previously been shown to suppress T helper 17 (Th17) polarization and cytokine secretion. Therefore, they are thought to be implicated in a fetoprotective role in vivo (*51*) since Th17 profiles lead to pregnancy complications (*32*). PSGs can be found in the serum of pregnant women and are essential for healthy pregnancies (*52*, *53*), where lower concentrations can be a sign of fetal growth restriction and preeclampsia.

We observed significant T, B, and NK cell phenotype and activation changes in response to trophoblast subtype secretome. However, we also observed notable differences in immune cell activation when comparing 2D and 3D culture environments, especially in PBMC exposure to CT TCM (Fig. 9). PBMCs incubated with CT TCM had lower inflammatory cytokine secretion compared to ST and EVT phenotypes, which also generally correlated to lower expression of activation markers on T and B cells (Fig. 4, 6-8, S3, S11, S13, S16). Although we saw many immunomodulatory surface markers expressed the highest in ST differentiation conditions (Fig. 3), ST TCM generally promoted higher inflammatory cytokine secretion in PBMCs, as well as higher activation markers in the cytokine secreting NK phenotype (Fig. 4-8, S3, S8, S11, S13, S16). Though many of these molecules have soluble forms that could be present in our TCM, this may indicate that the presentation of these molecules by STs has a greater immunosuppressive effect than ST-secreted factors. Additionally, higher levels of type 1 cytokines are associated with RSA and preterm delivery, whereas higher type 2 cytokines are associated with successful pregnancy (*54–56*). Interestingly, we saw higher secretion of both type 1 and type 2 cytokines from PBMCs cultured with ST TCM compared to CT and EVT TCM or positive controls (Fig. 4, S3). Further, the cytokine secretion profile from PBMCs cultured with EVT TCM is associated with vascularization and remodeling, a primary function of EVTs in vivo, and these cytokines (MCP-1, IL8) were significantly higher in 3D groups relative to 2D.

**Fig. 9.**
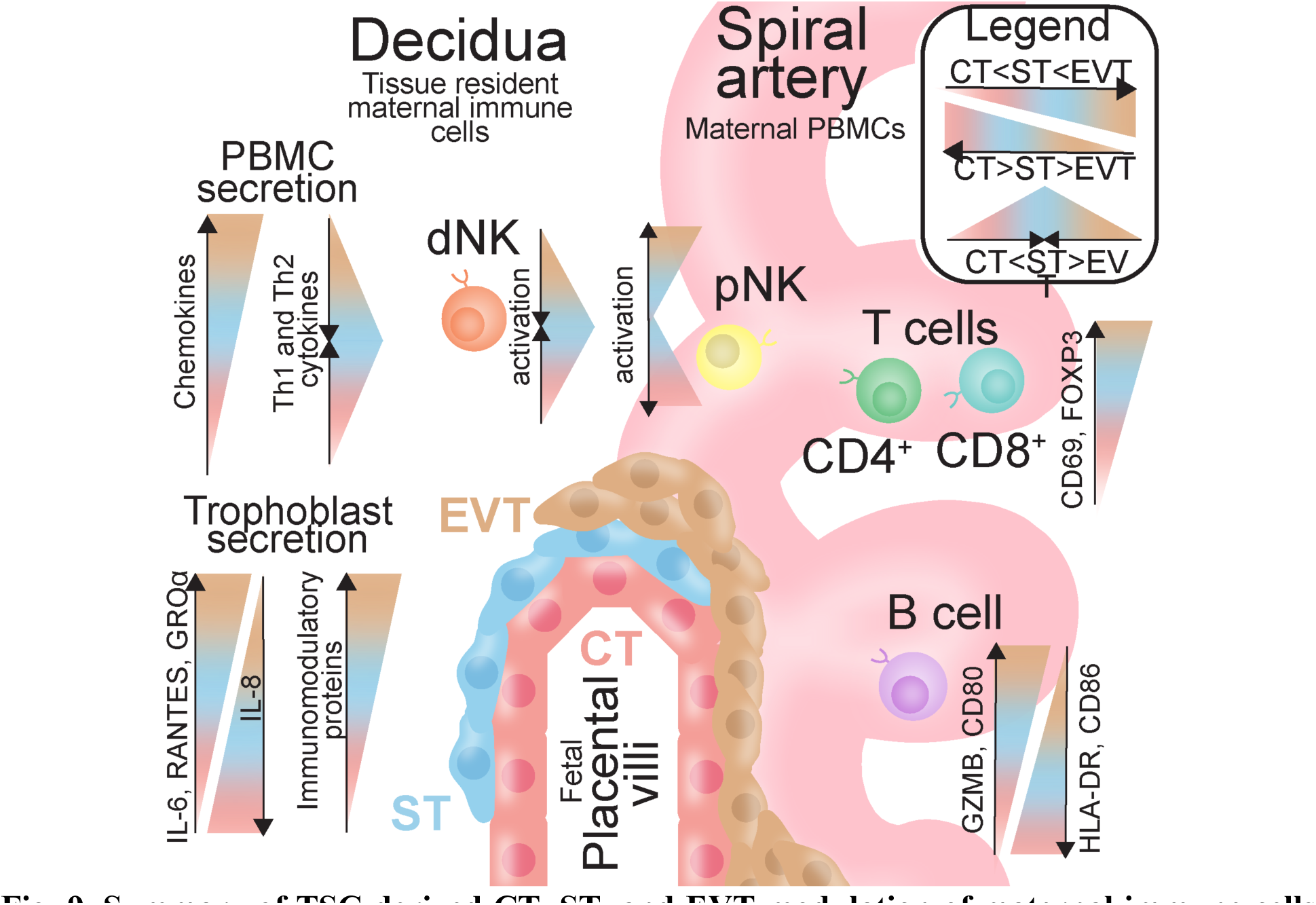
Summary of TSC-derived CT, ST, and EVT modulation of maternal immune cells. Placental trophoblast structure typically differentiates from CTs to STs and EVTs, with ST and EVT phenotypes having the greatest contact with maternal immune cells, whether in the decidual stroma or through the blood in the remodeling spiral arteries. Colored bars illustrate the amount of measured or expressed proteins identified in this study, with greater width denoting greater expression—pink, blue, and tan colors denote the CT, ST, and EVT phenotypes, respectively. Select markers are shown here to illustrate the increasing or decreasing marker or protein expression gradient from CT to ST to EVT.

Trophoblasts lack major histocompatibility complex (MHC) class II antigens and the MHC class I antigens HLA-A and -B; however, EVTs that come into contact with the maternal immune system express MHC class I antigens HLA-C, -E, and -G (*57–60*), which allow them to interact with killer-cell inhibitory receptors (KIR) on the surface of NK cells (*61*). These molecules are thought to be heavily involved in regulating NK cells at the maternal interface (*11*). CD16^+^ and CD56^++^ NK cells in unactivated PBMC conditions showed significantly higher expression of inhibitory receptor CD158a (KIR2DL1), which recognizes HLA-C (*62*), when cultured with EVT TCM compared to ST TCM (Fig. S8C, S8K). Conversely, degranulation marker CD107a (LAMP-1) was highest in CD16^+^ CD56^dim^ cytotoxic NK cells with ST TCM within unactivated PBMCs, whereas under activation conditions, CD107a had the highest expression with CT and EVT TCM (Fig. 5E, S8B, S8F). Although uterine cell frequencies of NK subtypes vary over time during pregnancy, peripheral blood NK cell frequencies remain unaffected (*61*), and we also did not see significant changes in CD16^+^ versus CD56^++^ frequencies in unactivated or activated PBMC populations with the addition of TCM (Fig. S7U-V). CD16^+^ and CD56^++^ NK cells displayed increased CD38 expression in unactivated PBMCs cultured in EVT TCM compared to CT (unactivated only) and ST TCM and 2D CT TCM compared to 3D CT TCM (Fig. S8D, S8L). Of note, the activated (CD38^+^) cytokine-secreting NK cell phenotype is associated with immune suppression, correlating to higher levels of circulating Tregs and poorer survival in patients with cancer (*63*). Interestingly, 2D CT TCM promoted higher expression of CD38 on both cytotoxic T and T helper cells in unactivated PBMCs compared to 3D CT TCM (Fig. S11D, S13D); however, in activated PBMCs, 2D CT TCM caused lower expression of CD38 on cytotoxic T and T helper cells compared to 3D CT TCM (Fig. S11J, S13J). CT and EVT trophoblast subsets may control and promote the increased CD38 expression in these immune cells in the peripheral blood of pregnant women compared to nonpregnant controls (*43*, *64*).

CD154 expression on T helper cells and soluble CD154 in serum of healthy pregnancy are significantly lower than in non-pregnant people and pre-eclamptic sera (*65*). Interestingly, in activated PBMCs, CD154 MFI was upregulated in T helper cells by TCM, with EVT TCM causing the highest CD154 expression; however, CT TCM showed a significantly lower frequency of CD154^+^ T helper cells compared to ST, EVT, and positive controls (Fig. S10K, S11I). CD40 expression in B cells, the cognate receptor to CD154 on T cells, was significantly lower in EVT and CT TCM conditions in unactivated and activated PBMC, respectively (Fig. S16G, S16N); therefore, the secretome of these cells may play a role in CD40 and CD154 expression in pregnant peripheral blood cells. Elevated levels of Granzyme B^+^ T cells and decreased Granzyme B^+^ NK cells are indicators of implantation failure (*66*). In unactivated PBMCs, T helper cells and B cells cultured in CT TCM had lower expression of Granzyme B than ST TCM, EVT TCM, and positive controls. However, in activated PBMCs, only 2D CT TCM lowered Granzyme B expression compared to 3D CT TCM (Fig. S11A, S11G, S16A, S16H). Interestingly, CT TCM also caused lower cytokine secretion from PBMCs compared to ST and EVT TCM in unactivated and activated conditions, corresponding to generally lower CD8^+^ cytotoxic T cell activation relative to other groups. Notably, when cultured in CT TCM, cytotoxic T cells had lower Granzyme B expression in both unactivated and activated PBMCs and lower CD25 and CD69 expression in activated PBMC conditions (Fig. 4, S13A, S13G-H, S13L). These modulations by CT TCM suggest CT may play a role in downregulating CD154 and modulating Granzyme B expression in CD4^+^ T cells and CD8^+^ cytotoxicity in the maternal periphery.

CT 3D TCM caused increased CD86 MFI in B cells in both unactivated and activated conditions compared to CT 2D TCM, which was decreased compared to activated controls and aligned with EVT TCM in the unactivated conditions (Fig. S16C, S16J). HLA-DR expression of B cells revealed the opposite trend, with higher expression resulting from culture with 2D compared to 3D CT TCM (Fig. S16B, S16I). These trends matched observations of HLA-DR expression in vivo, which is increased in the peripheral blood throughout pregnancy, except in the cases of intrauterine growth retardation, preeclampsia, and placental abruption (*67*). Interestingly, IL-10 secretion increased when PBMCs were cultured in ST TCM, which corresponded to higher CD71 MFI in B cells (Fig. S3H, S3U, S16D, S16K), which is characteristic of memory B cells (*68*). EVT TCM increased FOXP3 T cell expression, indicating an expanded Treg population, in unactivated PBMCs, and regulatory B cell markers CD24 and Granzyme B were also significantly increased compared to CT and ST (Granzyme B only) TCM (Fig. S11E, S13E, S16A, S16F). Overall, CT, ST, and EVT TCM modulated PBMC phenotypic expression comparable to healthy pregnancy-induced expression in the peripheral blood cells of pregnant people.

Our model has some limitations that may restrict the application of these results. Due to the small sample size of human PBMC samples tested, these donor-specific responses may only be widely applicable to a portion of the general population. Additionally, there is no information on whether the PBMC samples were from people who were previously pregnant or had pregnancy complications, which could significantly impact cell expression outcomes. Further, in this model, we assessed immune cell populations within whole PBMCs exposed to the TCM of TO containing predominantly one trophoblast phenotype. As such, the phenotypic shifts observed here may not be comparable to interactions involving trophoblast-immune cell contact and may not be generalizable to the secretome of a mixture of trophoblast phenotypes. Furthermore, this model may not be directly comparable to immune modulation at the maternal-fetal interface, as PBMCs are likely to react differently than tissue-resident immune cells. For example, Tregs can only be induced by P4 from cord blood cells, not PBMCs (*32*). Finally, media controls from CT, ST, and EVT cultures often produced significant immune cell phenotypic shifts due to growth factors and inhibitors; for example, CT and EVT media contain A83-01, and CT media contains SB431542 (TGFβ kinase inhibitors), which might explain the high abundance of TGFB1 expressed by STs compared to CTs and EVTs (Fig. 3O). Future studies investigating TO immunomodulation will aim to address the limitations listed here.

Here, we have developed an in vitro human placental organoid model to mechanistically evaluate the contributions of trophoblast phenotype and trophoblast culture microenvironment to maternal immune dynamics during human pregnancy. We assessed the effects of trophoblast culture condition on the trophoblast secretome and immunomodulatory protein expression, finding a phenotype-dependent cytokine and chemokine profile. Additionally, we assessed changes in phenotypic and activation markers of human T, B, and NK cells in response to trophoblast-secreted factors. We found that the CT phenotype exhibited the most immunosuppressive secretome across nearly all conditions. Furthermore, culture conditions significantly altered trophoblast secretome and immune cell responses, most commonly in the CT phenotype. Our results suggest that 3D synthetic hydrogel cultured placental organoids interact with immune cells more comparably to in vivo human sera than traditional 2D trophoblast culture.

## Materials and Methods

### Experimental Design

This study aimed to evaluate the effects of trophoblast phenotypes and trophoblast culture condition (2D versus 3D) on PBMCs during pregnancy using an in vitro early gestation human trophoblast organoid model. The immunomodulatory protein expression and secretion by trophoblast phenotypes were assessed using proteomics analysis and multiplex cytokine and chemokine secretion assays. Next, we evaluated trophoblast secretion from phenotypes and culture conditions on the peripheral blood cells of healthy female donors to assess phenotype changes caused by trophoblasts.

### Cell culture

All cells were maintained in a humidified incubator at 37 °C in 5% CO_2_. Human TSC CT 1049, derived from 6-week gestation age placental tissues (*20*), were kindly gifted by Dr. Mana Parast from the University of California, San Diego. These cells (CT phenotype) were cultured with modifications to previously published literature (*18*) on 5 µg/mL collagen-coated (Sigma catalog #C0543-1VL) 6-well plates in advanced DMEM/F12 (Gibco^TM^ catalog #12634010), 1X B27 (Gibco catalog #17504044), 1X N2 (Gibco catalog #17502048), 1X GlutaMAX (Gibco catalog #35050061), 150 µM 1-thioglycerol (Sigma catalog #M6145), 1% KnockOut Serum Replacement (Gibco catalog #10-828-028), 0.05% BSA (Gemini Bio Products catalog #50-753-3079), 2 µM CHIR99021 (Sigma catalog #SML1046), 500 nM A83-01 (Tocris Bioscience^TM^ catalog #29-391-0), 1 µM SB43154 (Sigma catalog #616464), 0.8 mM valproic acid sodium salt (Sigma catalog #676380), 5 µM Y27632 (Selleck catalog #S1049), 100 ng/mL FGF2 (R&D Systems #3718FB01M), 50 ng/mL EGF (R&D Systems catalog #236-EG-01M), 50 ng/mL HGF (Stem Cell Technologies catalog #78019), and 20 ng/mL Noggin (R&D Systems catalog #6057-NG) with media replenished every 2-3 days or passaged with TrypLE (Gibco catalog #1260421).

ST differentiation was achieved by culturing passaged CT cells on 5 ug/mL collagen-coated 6- or 24-well plates (2D conditions only) in advanced DMEM/F12, 1X B27, 1X N2, 4% KnockOut Serum Replacement, 0.05% BSA, 2.5 mM Y27632, and 2 mM forskolin (Sigma catalog #344270) with media replenished every 2 days.

EVT differentiation was achieved by culturing passaged CT cells on 20 µg/mL fibronectin (Gibco catalog #PHE0023, incubated >90 minutes in DPBS at 37 °C) coated 6- or 24-well plates (2D conditions only) in advanced DMEM/F12, 0.1 mM 2-mercaptoethanol, 0.3% BSA, 1% ITS-X supplement (Gibco catalog #51500-056), 7.5 µM A83-01, 4% KnockOut Serum Replacement, 100 ng/mL NRG1 (Cell Signaling catalog #26941),, and 2% Matrigel (Corning catalog #354234) for 4 days (changed every 2 days), then changing the media to 0.5% Matrigel without NRG1 for another 2 days.

The Jurkat (clone E6-1) cell line was purchased from ATCC (TIB-152 ^TM^) and cultured according to ATCC’s handling information. Briefly, these cells were cultured in RPMI 1640 (Gibco catalog #11875119) with 10% FBS (Gibco catalog #26140095), 2 mM L-glutamine (Gibco catalog #25030081), 10 mM HEPES (Gibco catalog #15630130), 1 mM sodium pyruvate (Gibco catalog #11360070), and 1X non-essential amino acids (NEAA, Gibco catalog #11140076) and maintained at a cell concentration between 10^5^ to 10^6^ viable cells per mL with media changes every 2-3 days.

The NK-92 cell line was purchased from ATCC (CRL-2407) and cultured in Alpha MEM without ribonucleosides (Gibco catalog #12000-063), 12.5% FBS, 12.5% Horse Serum (Gibco catalog #16050122), 1.5 g/L sodium bicarbonate (Sigma catalog #S5761), 0.2 mM Myo-inositol (Sigma catalog#I-7508), 0.1 mM 2-mercaptoethanol (Gibco catalog#21985-023), 0.02 mM folic acid (Sigma catalog#F-8758), and 100 U/mL premium grade recombinant human IL-2 (Sigma catalog #I7908) with media changes every 2-3 days to maintain the cells at a concentration of 0.2 M cells/mL.

PBMCs from female donors (32-45 years old, white or Caucasian, non-smokers, BMI 19.92-26.63, CMV negative) were purchased from BioIVT and stored frozen in CryoStor CS10 in the vapor phase of (LN_2_) until use. PBMCs were rested in RPMI + 10% FBS for ∼24 hours before use.

### Hydrogel preparation

CT cells were grown in traditional 2D conditions in CT media until 60-80% confluence and passaged with TrypLE. CT cells (100, 000) were encapsulated in a 15 µL hydrogel for 6 days maintained in CT media, or 200,000 CT cells were encapsulated in the same volume and cultured in ST or EVT differentiation medium. PEG-mal (20 kDa, 4-arm, 5% w/v, Laysan Bio catalog #4arm-PEG-MAL-20K) macromer with 1 mM adhesion ligand (GFOGER, GGYGGGPP(GPP)5GFOGER(GPP)5GPC, GenScript) and protease-degradable VPM (CRDVPMSMRGGDRCG, GenScript) (*69–71*) or nondegradable DTT (Thermo Scientific^TM^ #R0861) were used to encapsulate cells as previously described (*72*, *73*). Briefly, PEG-mal, adhesion ligands, and crosslinker reagents were resuspended in DPBS, and the pH was adjusted to 7 if necessary, using sodium hydroxide. Matrigel (Corning catalog #356237) was diluted to 5 mg/mL (0.5% w/v) with cell-specific media, and 2D conditions were plated with the same number of cells per subtype condition into a 24-well culture-treated plate following coating methods above. Cell-specific media (1 mL) was added to the hydrogels or 2D wells and replaced every 2 days based on the culture procedure above. Day 6 trophoblast conditioned media (TCM) was collected from the conditions, spun down to remove cell debris, and frozen at -80 °C until use.

### TSC secretion assessment

CTs were encapsulated in PEG-GFOGER-DTT, PEG-GFOGER-VPM, or Matrigel or cultured in 2D for 6 days with CT, ST, or EVT differentiation media as stated above, with media changes on days 2 and 4. Supernatants were collected on day 6 and immediately assessed for cytokine and chemokine secretion by LEGENDplex™ using LEGENDplex Human Inflammation Panel 1 (13-plex) with V-bottom Plate (Biolegend catalog #740809; IL-1β, IFN-α2, IFN-γ, TNF-α, MCP-1, IL-6, CXCL8 (IL-8), IL-10, IL-12p70, IL-17A, IL-18, IL-23, IL-33) and LEGENDplex HU Proinflam. Chemokine Panel 1 (13-plex) w/VbP (Biolegend catalog #740985; CXCL8 (IL-8), CXCL10 (IP-10), CCL11 (Eotaxin), CCL17 (TARC), CCL2 (MCP-1), CCL5 (RANTES), CCL3 (MIP-1α), CXCL9 (MIG), CXCL5 (ENA-78), CCL20 (MIP-3α), CXCL1 (GROα), CXCL11 (I-TAC), CCL4 (MIP-1β)) according to the manufacturer’s instructions. LEGENDplex data was analyzed using the LEGENDplex Data Analysis Software Suite from Qognit.

### Proteomics

CTs were encapsulated in degradable synthetic (PEG-VPM-GFOGER) and natural (Matrigel) hydrogels or 2D-cultured for 6 days with media changes every 2 days for CT, ST, and EVT differentiation conditions as detailed above. On day 6, 2D cells were washed with DPBS and lifted with TrypLE, and hydrogels were washed in DPBS for 10 minutes to remove serum. Cell pellets and hydrogels were snap-frozen in liquid nitrogen for lysis and stored at -80 °C until sample preparation and analysis. Proteomics sample preparation was done as described (*21*). Briefly, samples were lysed in 8M urea lysis buffer, and the protein concentration of clarified lysate was determined using BCA (bicinchoninic acid) Protein Assay (Pierce). An equal amount of protein per sample was digested by trypsin (Promega), and the resulting peptides were quantified by BCA assay. Proteomics data were acquired on Ultimate U3000 RSLCnano liquid chromatography system coupled to a Thermo Orbitrap Eclipse mass spectrometer equipped with a FAIMS Pro interface (ThermoFisher Scientific) using Data Independent Acquisition (DIA) - Mass Spectrometry (MS). DIA-MS data was queried against a sample-specific spectral library (generated by offline fractionation of a pooled reference sample) in Spectronaut using default settings with cross-run normalization turned “OFF,” and data imputation was disabled. Results were filtered to 1% protein and peptide FDR in Spectronaut.

Protein abundances from Spectronaut were normalized with variance stabilizing normalization in R (v4.2.1). Normalized protein abundances were log2-transformed and used for heatmap and principal component analysis (PCA) in R (v4.4.0). Heatmaps employed hierarchical clustering of both samples and proteins based on Euclidean distance. Relevant proteins are only shown on heatmaps if present in more than 50% of the samples. For PCA plots, missing values were set to zero.

### Cell culture with TCM

NK-92, Jurkats, or PBMCs were plated at 100, 000 cells in 200 µL of TCM, positive controls, or media controls. TCM from day 6 CT, ST, and EVT differentiation conditions were used, along with PEG-GFOGER-DTT, PEG-GFOGER-VPM, Matrigel, and 2D cultured condition cells. Positive controls used were TGFβ1 (NK-92: 10 ng/mL; Jurkat and PBMC: 50 ng/mL, R & D Systems catalog #1064-ILB-010), TGFβ1 and IL-10 (20 ng/mL; BioLegend catalog #571004), progesterone (P4, 4 µg/mL, Sigma catalog #P8783-5G), human chorionic gonadotropin (hCG, for Jurkat: high 1000 IU/mL and low 200 IU/mL; PBMC: 500 IU/mL, Sigma catalog #CG5-1VL). Media controls were used to determine if media components alone affect protein secretion and expression. Hence, these controls were CT, ST, EVT, or cell-specific media (NK-92, Jurkat, or PBMC media as outlined above) conditions without cell secretions. For unactivated conditions, NK-92 and PBMCs used to assess natural killer (NK) cell populations were cultured for 26 hours in TCM or controls, whereas Jurkats and PBMCs used to determine T and B cell populations or inflammatory cytokine secretions were cultured for 48 hours. For activated conditions, cells were cultured for 24 hours in TCM or controls, then cultured with added 10 ng/mL phorbol 12-myristate 13-acetate (PMA, Sigma catalog #524400) and 1 ug/mL ionomycin calcium salt (ionomycin, Sigma catalog #I3909) to TCM or controls for 2 hours (NK-92 and PBMCs used to assess NK markers) or 24 hours (PBMC secretion data, Jurkats, and PBMCs used assess T and B cell protein expression). In the last 4 hours of cell culture, 1X brefeldin A (BioLegend catalog #420601) was added to conditions to detect Granzyme B or IFNγ if necessary. Supernatants were collected at these time points (NK-92: 26 total hours, PBMCs: 48 total hours) to assess cell secretion (NK-92: by enzyme-linked immunosorbent assay (ELISA) as outlined below, PBMC: by LEGENDplex as outlined below) and cells were stained for flow cytometry according to the methods below.

### NK-92 IFNγ ELISA

NK-92 supernatants were collected after a 26-hour incubation with TCM with or without adding 10 ng/mL PMA and 1 ug/mL ionomycin at the 24-hour mark. Supernatants were spun down to remove cell debris and frozen at -80 °C until analysis using the ELISA MAX™ Deluxe Set Human IFN-γ (BioLegend catalog #430104) according to the manufacturer’s instructions. A four-parameter logistic curve was used to determine concentrations using MyAssays.com.

### PBMC secretion assessment

Supernatants were collected after a 48-hour PBMCs and TCM incubation and immediately assessed for cytokine secretion using LEGENDplex Human Inflammation Panel 1 (Biolegend catalog #740809) according to the manufacturer’s instructions. LEGENDplex data was analyzed using the LEGENDplex Data Analysis Software Suite from Qognit.

### Flow cytometry

Cells were collected and washed with PBS with 1% BSA and 2 mM EDTA. Dead cells, stained with Zombie Aqua^TM^ Fixable Viability Kit (BioLegend catalog #423102) at 1:1000 in DPBS for 30 minutes at room temperature following the manufacturer’s instructions, then washed with DPBS with 1% BSA, were gated out in all experiments. Cells were incubated with Human TruStain FcX^TM^ (FC Receptor Blocking Solution, 1:250, BioLegend catalog #422302) for 5 minutes at room temperature at a 1:250 dilution in DPBS with 1% BSA. Brilliant Stain Buffer PLUS (BD Horizon^TM^ catalog #BD566385) was then added into the staining cocktails as outlined below according to the manufacturer’s instructions, and stains were incubated with the cells for 15-20 minutes at room temperature in the dark. Cells were fixed and permeabilized for intracellular staining of Granzyme B and FOXP3 using FluoroFix Buffer and Intracellular Staining Permeabilization Buffer (Biolegend catalog #421002) according to the manufacturer’s instructions. FMO control samples were made by mixing a portion of each sample. Cells were fixed with FluoroFix^TM^ Buffer according to the manufacturer’s instructions (BioLegend catalog #422101) for 1 hour at 4°C or until analysis on a Life Technologies Attune NxT with Autosampler. UltraComp eBeads Plus (Invitrogen catalog #01-222-42) and ArC™ Amine Reactive Compensation Bead Kit (Invitrogen catalog #A10346) were stained as single-color controls for compensation. Samples were analyzed in FlowJo^TM^ 10.

The following anti-human antibodies were used in the Jurkat staining panel: anti-CD4 FITC (BioLegend catalog #317408, clone OKT4), anti-CD25 PerCP/Cyanine (Cy) 5.5 (BioLegend catalog #356112, clone M-A251), anti-HLA-DR APC (BioLegend catalog #327022, clone LN3), anti-CD38 APC/Cy7 (BioLegend catalog #397118, clone S17015A), anti-CD3 Brilliant Violet (BV) 421^TM^ (BioLegend catalog #317344, clone OKT3), anti-CD71 BV650 ^TM^ (BioLegend catalog #334116, clone CY1G4), anti-CD40L (CD154) BV 785 ^TM^ (BioLegend catalog #310842, clone 24-31), anti-CD69 PE/Dazzle ^TM^ 594 (BioLegend catalog #310942, clone FN50).

The following anti-human antibodies were used in the NK-92 staining panel: anti-CD107a (LAMP-1) FITC (BioLegend catalog #328606, clone H4A3), anti-CD56 (NCAM) BV650 ^TM^ (BioLegend catalog #362532, clone 5.1H11), and anti-CD16 BV785 ^TM^ (BioLegend catalog #302046, clone 3G8).

The following anti-human antibodies were used in the PBMC staining panel for purity: anti-CD4 FITC (BioLegend catalog #317408, clone OKT4), anti-CD20 PerCP/Cy5.5 (BioLegend catalog #375510, clone S18015E), anti-CD34 APC (BioLegend catalog #343510, clone 581), anti-CD3 APC/Cy7 (BioLegend catalog #344818, cloneSK7), anti-CD45 421^TM^ (BioLegend catalog #304032, clone HI30), anti-CD8 BV650^TM^ (BioLegend catalog #344730, clone SK1), anti-CD16 BV785^TM^ (BioLegend catalog #302046, clone 3G8), anti-CD56 PE/Dazzle ^TM^ 594 (BioLegend catalog #362544, clone 5.1H11), and anti-CD14 PE/Cy7 (BioLegend catalog #301814, clone M5E2).

The following anti-human antibodies were used in the PBMC staining panel for B cells: anti-CD19 Alexa Fluor® 488 (BioLegend catalog #302219, clone HIB19), anti-CD3 PerCP/Cy5.5 (TONBO catalog #65-0037-T100, clone OKT3), anti-HLA-DR APC (BioLegend catalog #327022, clone LN3), anti-Granzyme B APC/Fire^TM^ 750 (BioLegend catalog #372210, clone QA16A02), anti-CD86 BV421 (BioLegend catalog #305426, clone IT2.2), anti-CD71 BV650 (BioLegend catalog #334116, clone CY1G4), anti-CD80 BV785 (BioLegend catalog #305238, clone 2D10), anti-CD24 PE/Dazzle 594 (BioLegend catalog #311134, clone ML5), and anti-CD40 PE/Cy7 (BioLegend catalog #334322, clone 5C3).

The following anti-human antibodies were used in the PBMC staining panel for NK cells: (donors 1-2) anti-IFN-γ FITC (BioLegend catalog #506504, clone B27), anti-CD19 PerCP/Cy5.5 (BioLegend catalog #309504, clone QA18A75), anti-Granzyme B APC (BioLegend catalog #303522, clone HIT2), anti-CD107a (LAMP-1) APC/Fire 750 (BioLegend catalog #328654, clone H4A3), anti-CD3 BV421 (BioLegend catalog #317344, clone OKT3), anti-CD56 (NCAM) BV650 (BioLegend catalog #362532, clone 5.1H11), anti-CD16 BV785 (BioLegend catalog #302046, clone 3G8), anti-CD158a (KIR2DL1) PE (BioLegend catalog #374904, clone HP-DM1), and anti-CD38 PE/Dazzle 594 (BioLegend catalog #303538, clone HIT2) or (donors 3-4) anti-CD19 AF488 (BioLegend catalog #302219, clone HIB19), anti-CD3 PerCP/Cy5.5 (TONBO catalog #65-0037-T100, clone OKT3), anti-Granzyme B APC (BioLegend catalog #303522, clone HIT2), anti-CD107a (LAMP-1) APC/Fire 750 (BioLegend catalog #328654, clone H4A3), anti-CD56 (NCAM) BV650 (BioLegend catalog #362532, clone 5.1H11), anti-CD16 BV785 (BioLegend catalog #302046, clone 3G8), anti-CD158a (KIR2DL1) PE (BioLegend catalog #374904, clone HP-DM1), and anti-CD38 PE/Dazzle 594 (BioLegend catalog #303538, clone HIT2).

The following anti-human antibodies were used in the PBMC staining panel for T cells: anti-CD4 FITC (BioLegend catalog #317408, clone OKT4), anti-CD26 PerCP/Cy5.5 (BioLegend catalog #356112, clone M-A251), anti-Granzyme B APC (BioLegend catalog #303522, clone HIT2), anti-CD3 APC/Cy7 (BioLegend catalog #344818, clone SK7), anti-CD8 BV650 (BioLegend catalog #344730, clone SK1), anti-CD40L (CD154) BV785 (BioLegend catalog #310842, clone 24-31), anti-CD38 PE (BioLegend catalog #303506, clone HIT2), anti-FOXP3 PE/Dazzle 594 (BioLegend catalog #320126, clone 206D), and anti-CD69 PE/Cy7 (BioLegend catalog #310912, clone FN50).

### Statistical analysis

Flow cytometry data was analyzed using FlowJo^TM^ version 10.8.1. MFI is shown as median fluorescence intensity and percent positive cells from the parent population. LEGENDplex data was analyzed using the LEGENDplex Data Analysis Software Suite from Qognit. LEGENDplex, ELISA, and flow cytometry data were formatted with Microsoft Excel, and graphs and statistics were completed using GraphPad Prism 10 version 10.2.1. Ordinary one- or two-way ANOVAs with Dunnett’s or Tukey’s multiple comparisons tests, as indicated in the figure caption, were used for statistical significance. Data are shown as individual hydrogel replicates with mean ± SEM unless otherwise stated. Proteomics data underwent analysis using an ANOVA followed by Tukey post hoc to correct for multiple hypothesis testing.

## Acknowledgments

We appreciate the Arizona State University KE core facility Flow Cytometry Core for providing the Attune NxT, and Adam Kindelin from this core for guidance during experimental planning. We also thank Dr. Mana Parast for kindly gifting the TSC CT 1049 cell line used in this article. Pipette artwork shown in Fig. S4 and S5 was used from Servier Medical Art (https://smart.servier.com/), licensed under a Creative Commons (CC) BY 4.0 DEED Attribution License. We thank ASU’s Regenerative Medicine and Bioimaging Facility for providing the Leica SP8 confocal microscope system, acquired by NIH SIG Award 1 S10 OD023691-01. This reported research includes work performed in the Integrated Mass Spectrometry Shared Resource supported by the National Cancer Institute (NCI) of the NIH under grant number P30CA033572.

## Author contributions

Conceptualization: EMS, JDW

Investigation: EMS, EMB, MWB, RS

Formal Analysis: EMS, EMB, MWB, NH, JDW

Writing – original draft: EMS, JDW

Writing – review & editing: EMS, EMB, MWB, NH, RS, PP, JDW

Funding Acquisition: JDW

## Data and materials availability

All data are in the main text or the supplementary materials.

