## Supplemental Figures for "Human placental organoids as a model to probe early gestation maternal immune dynamics"

**This PDF file includes:**

Figs. S1 to S16

**Other supporting information**

Supplementary Tables: Tables S1-S2

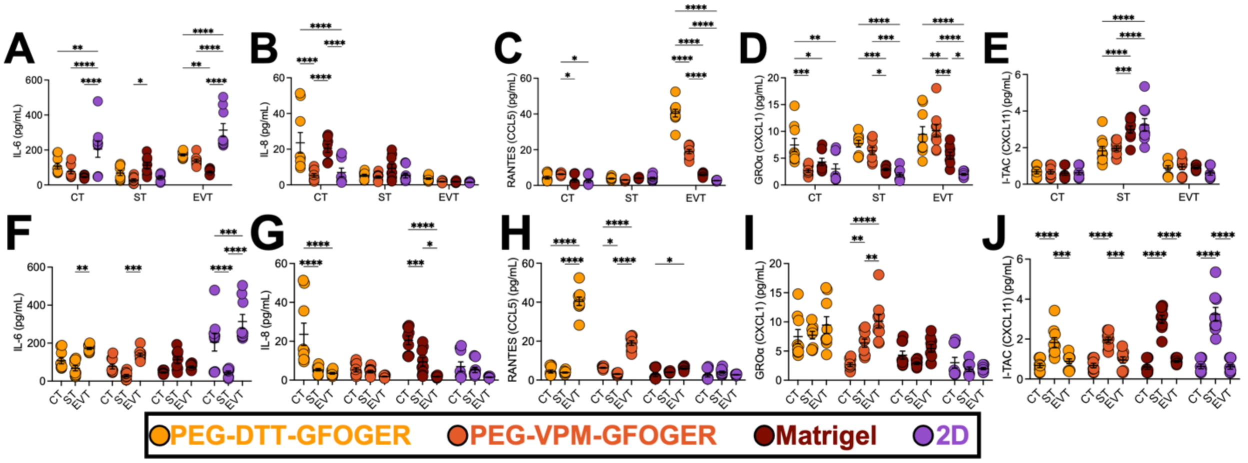


**Fig. S1. Alternate statistical analysis of cytokines and chemokines secreted from trophoblast secretome.** Secretion of (**A, F**) IL-6, (**B, G**) IL-8, (**C, H**) RANTES, (**D, I**) GROα, and (**E, J**) I-TAC from CT, ST, and EVT organoids on day 6 cultured in PEG-GFOGER-DTT, PEG-GFOGER-VPM, Matrigel, or 2D TSCs by LEGENDplex$^{TM}$. Data are shown as mean ± SEM and analyzed by ordinary two-way ANOVA with Tukey's multiple comparisons test (**A-E**) between culture conditions or (**F-J**) between CT, ST, and EVT differentiation conditions. * $p < 0.05$, ** $p < 0.01$, *** $p < 0.001$, **** $p < 0.0001$. n=9 from 3 independent experiments.

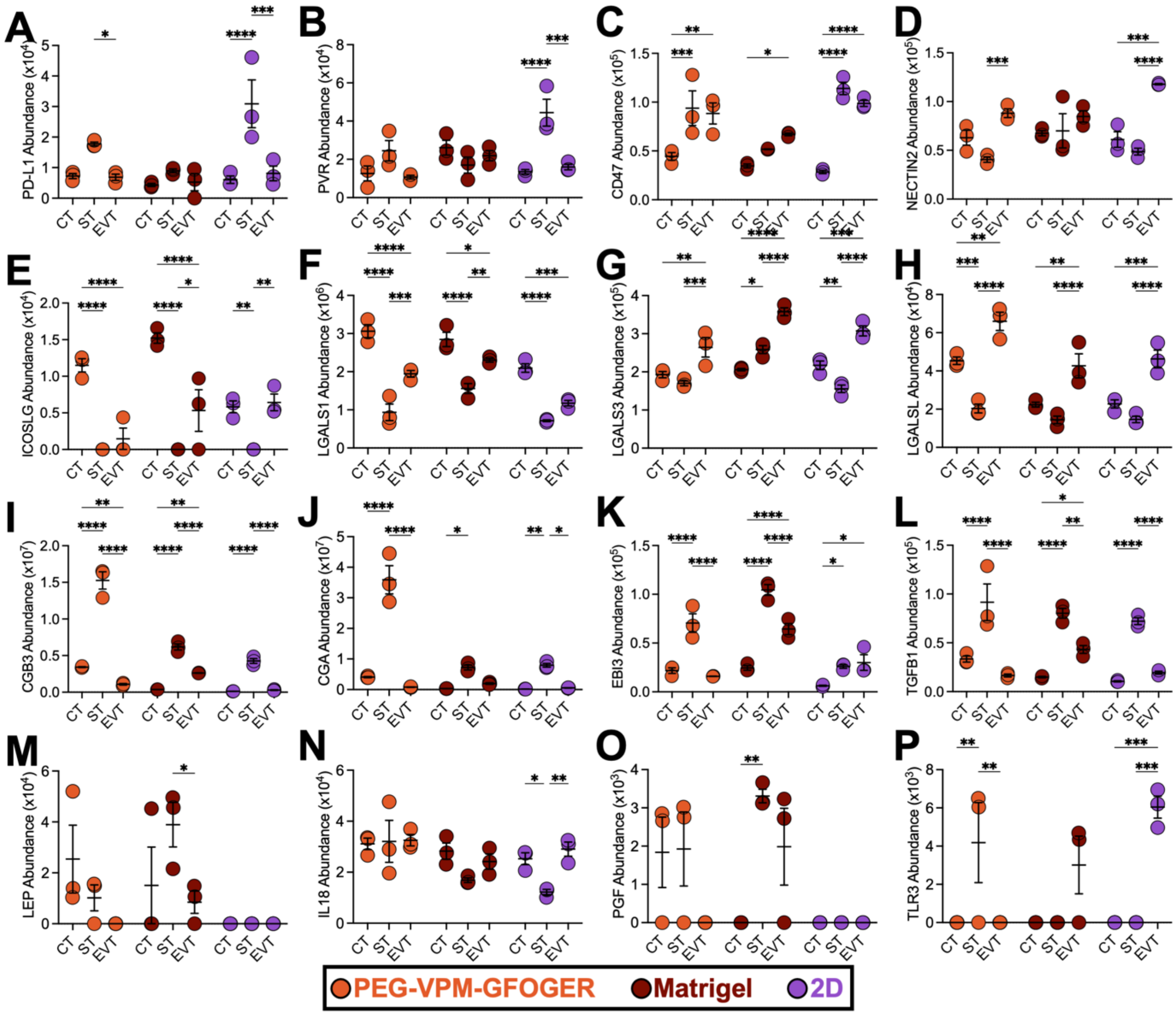

**Fig. S2. Trophoblast phenotype modulates the expression of immunomodulatory proteins.** (**A**) PD-L1, (**B**) PVR, (**C**) CD47, (**D**) NECTIN2, (**E**) ICOSLG, (**F**) LGALS1, (**G**) LGALS3, (**H**) LGALSL, (**I**) choriogonadotropin subunit beta 3 (CGB3), (**J**) CGA, (**K**) EBI3, (**L**) transforming growth factor beta-1 (TGFB1), (**M**) LEP, (**N**) IL-18, (**O**) placental growth factor (PGF), and (**P**) toll-like receptor (TLR) 3. Data are shown as mean ± SEM and analyzed by two-way ANOVA with Tukey's multiple comparisons test. * $p < 0.05$, ** $p < 0.01$, *** $p < 0.001$, **** $p < 0.0001$. n=3.

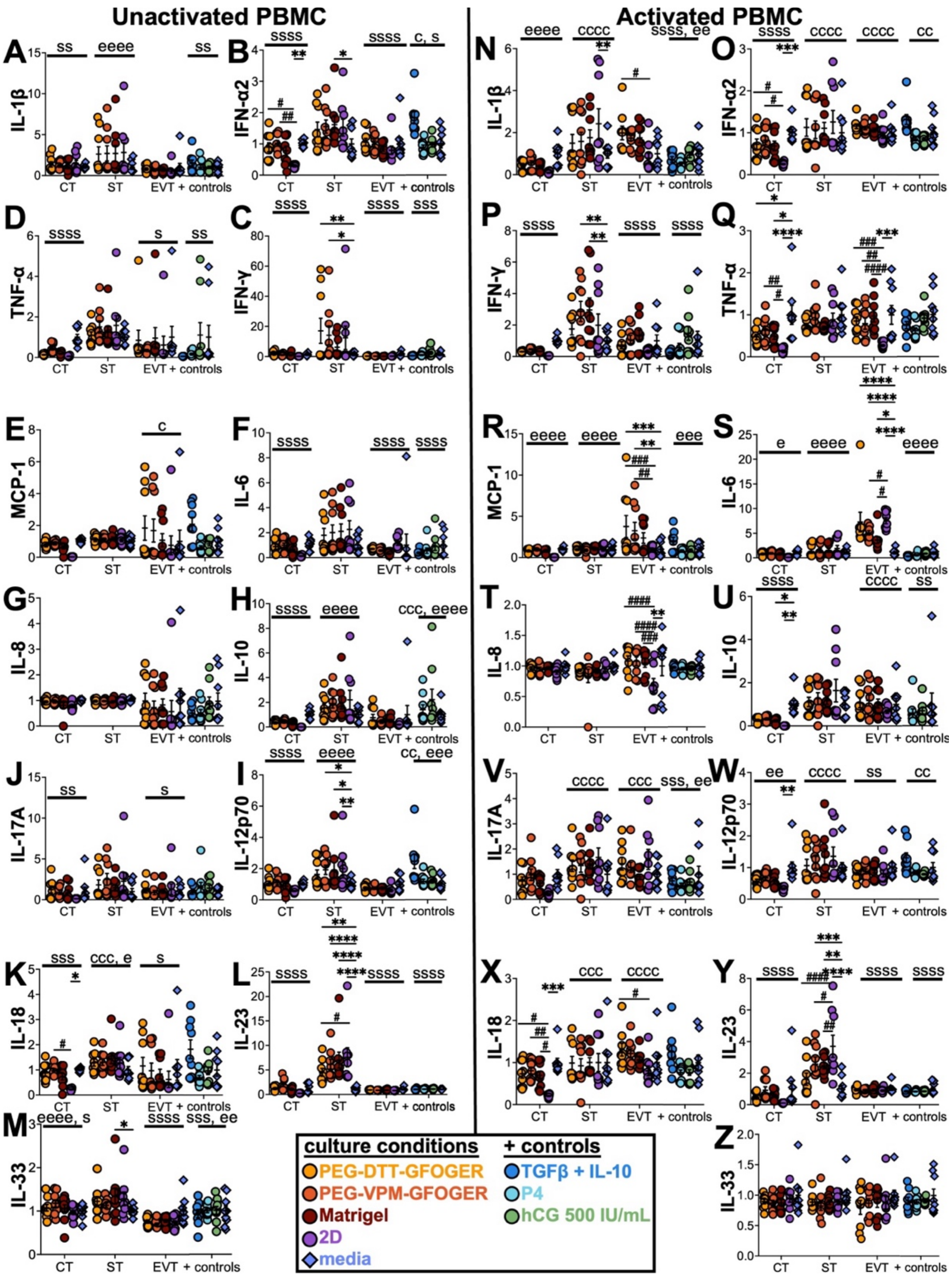


**Fig. S3. Cytokine secretion from PBMC with statistics compared to media or 2D controls.** (**A-M**) Normalized PBMC secretion from unactivated and (**N-Z**) activated with PMA and ionomycin conditions using the experimental layout in Fig. 4A. Data are shown as mean ± SEM and analyzed by ordinary two-way ANOVA with Dunnett's multiple comparisons test to the 2D (#) and media (*) controls or Tukey's multiple comparisons test between CT, ST, EVT, and positive (+) control conditions (significance to c, s, and e, respectively, media controls not included). * $p < 0.05$, ** $p < 0.01$, *** $p < 0.001$, **** $p < 0.0001$. n=9 from 3 independent experiments.

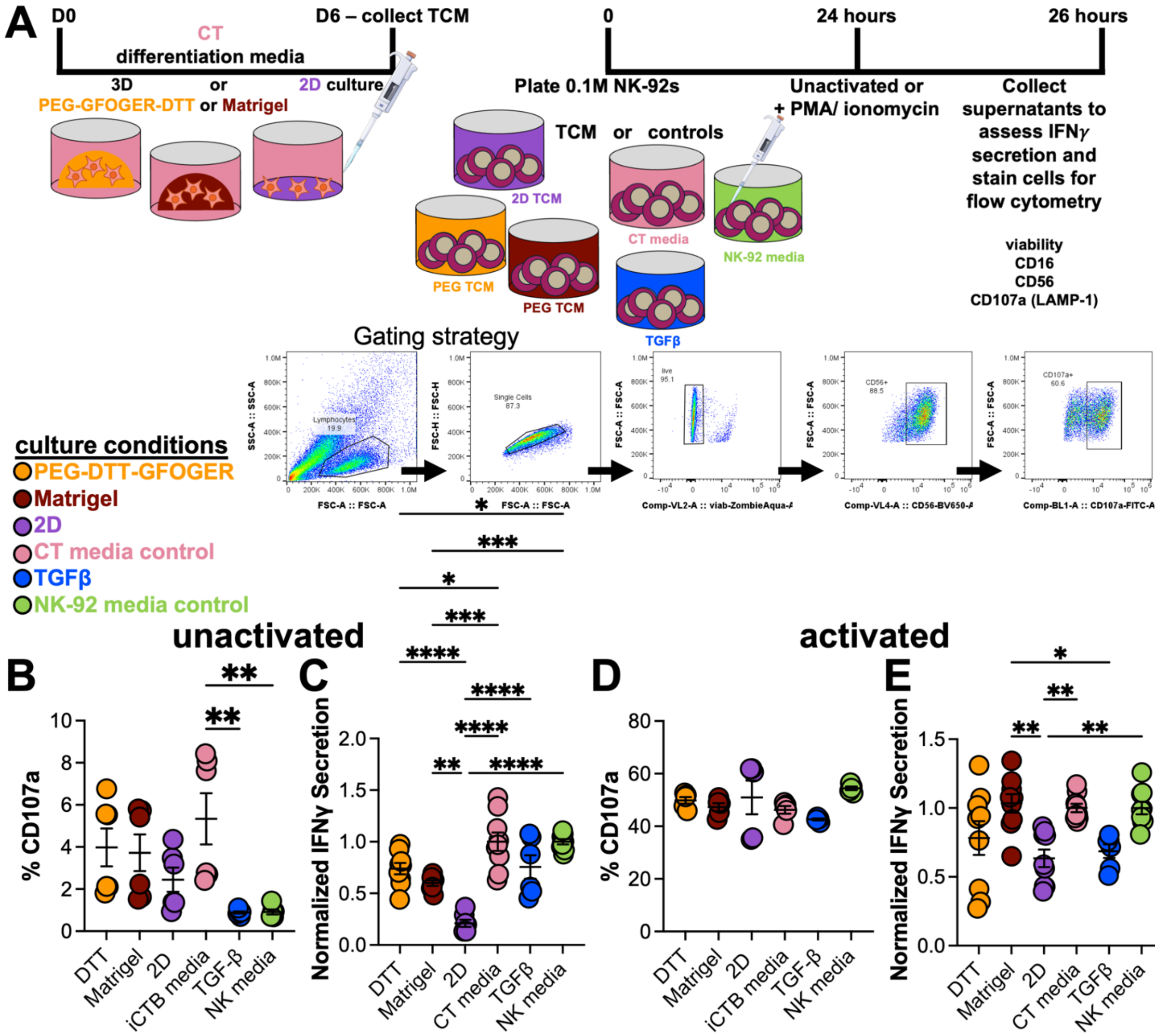


**Fig. S4. NK-92 cell line is modulated by CT secretome.** (**A**) Schematic of experimental layout. TCM was collected on day 6 from CT grown in PEG-GFOGER-DTT (DTT), Matrigel, or 2D. NK-92s were plated with TCM or controls (CT media, NK-92 media, and TGFβ) for 26 hours and left unactivated or activated with PMA and ionomycin at the 24-hour time point. Supernatants were collected and assessed for IFN-γ, and flow cytometry was run on the cells to assess the CD107a degranulation marker. (**B-C**) Unactivated or (**D-E**) activated NK-92 with TCM (**B, D**) percentage of CD107a$^{+}$ cells and (**C, E**) secretion of IFNγ normalized to respective media controls. Data are shown as mean ± SEM and analyzed by ordinary one-way ANOVA with Dunnett's multiple comparisons test to the 2D control. * $p < 0.05$, ** $p < 0.01$, *** $p < 0.001$, **** $p < 0.0001$. n=6-9 from 2-3 independent experiments.

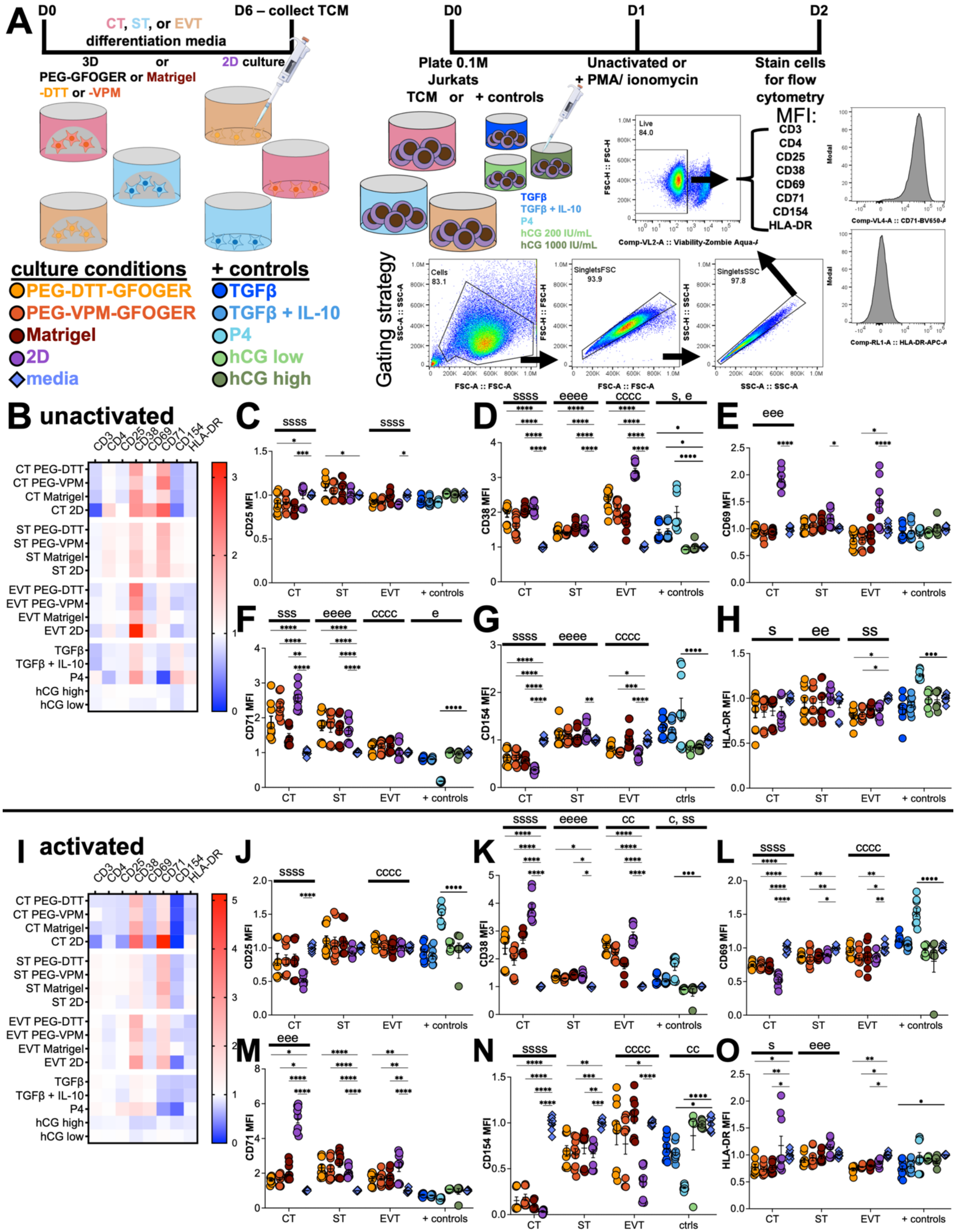


**Fig. S5. Jurkat protein markers are modulated by TCM.** (**A**) Schematic of experimental layout. TCM was collected on day 6 after CT, ST, or EVT culture conditions grown in PEG-GFOGER-DTT, PEG-GFOGER-VPM, Matrigel, or 2D. 100, 000 Jurkat cells were plated with TCM or controls for 2 days and left unactivated or activated with PMA and ionomycin at the 24-hour time point. Cells were stained and assessed by flow cytometry of common activation markers after 48 hours. Protein markers from (**B-H**) unactivated or (**I-O**) activated Jurkats incubated with TCM. (**B, I**) Overall heatmap of protein markers and (**C-H, J-O**) protein expression MFI normalized to condition-specific media controls – (**C, J**) CD25, (**D, K**) CD38, (**E, L**) CD69, (**F, M**) CD71, (**G, N**) CD154, and (**H, O**) HLA-DR normalized to media controls and standardized. Data are shown as mean ± SEM and analyzed by ordinary two-way ANOVA with Dunnett's multiple comparisons test to the 2D control (*) or main effects only and Tukey's multiple comparisons test between CT, ST, EVT, and + controls (c, s, e, respectively). * $p < 0.05$, ** $p < 0.01$, *** $p < 0.001$, **** $p < 0.0001$. n=9 from 3 independent experiments.

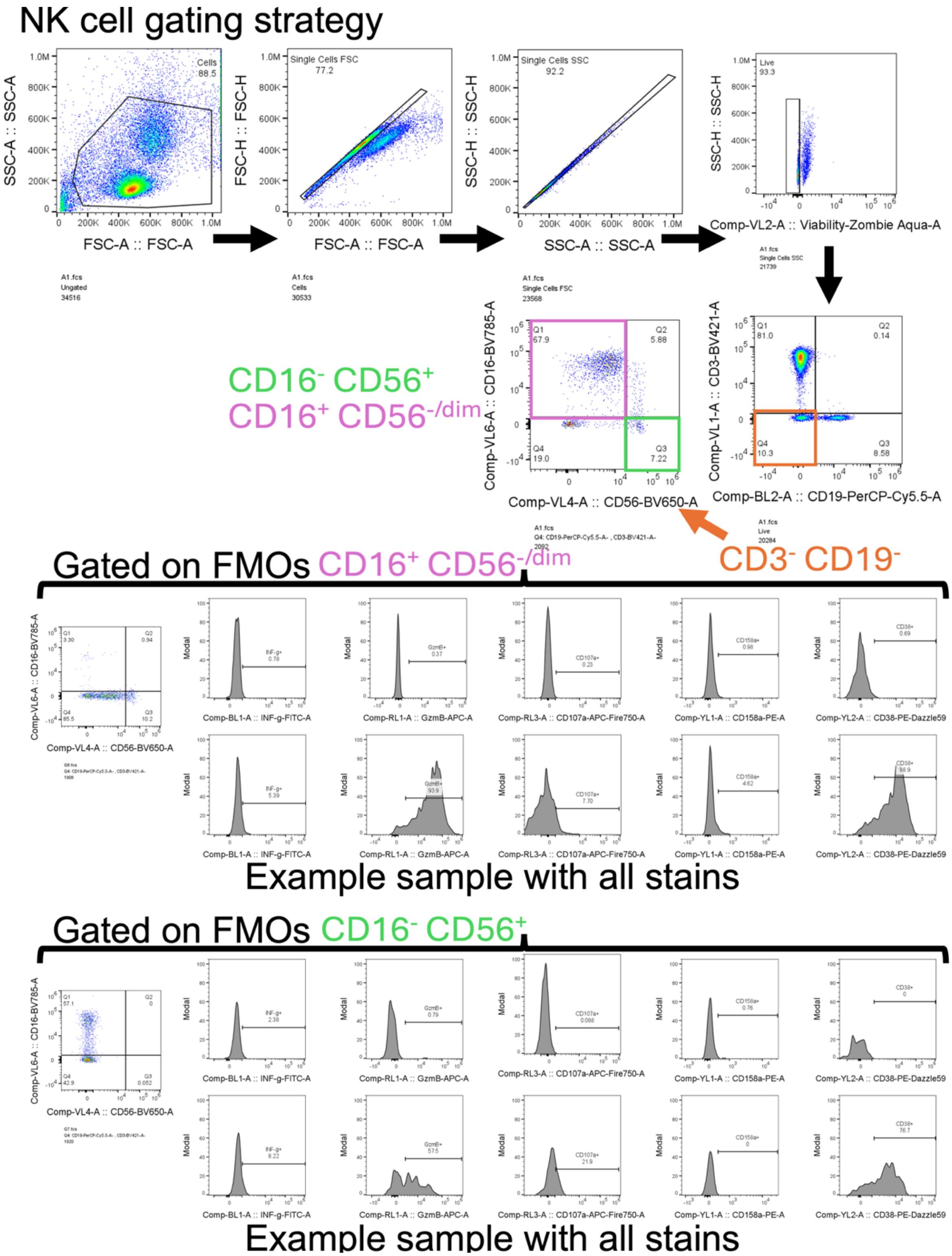


**Fig. S6. NK cell gating strategy from PBMC population.** Debris was gated out, singlets and live events were gated on, then $CD3^+$ and $CD19^+$ cells were also gated out. The $CD3^-$ $CD19^-$ population was assessed for $CD16^+$ and $CD56^+$ subsets to evaluate MFI and percentage of positive cells of protein markers, where fluorescence minus one (FMO) controls were used to gate out cells negative for markers.

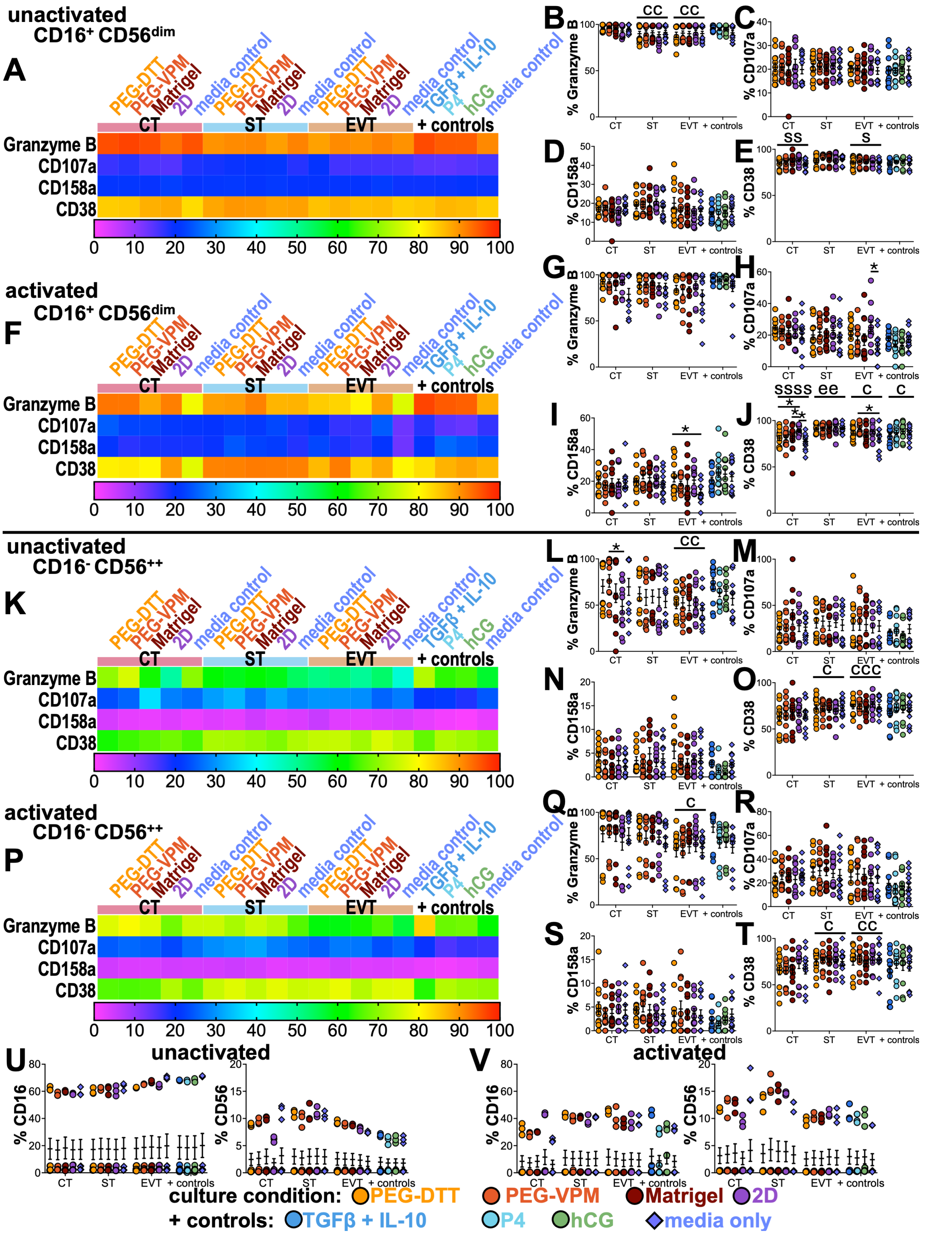


**Fig. S7. Trophoblast phenotype significantly affects the percentage of NK cells with activation markers within PBMCs.** (**A**) Heatmap of unactivated cytotoxic NK cell (CD16+ CD56dim) percent positive protein markers after a 26-hour culture of PBMC with TCM or positive

(+) controls evaluated by flow cytometry and percentages of (**B**) Granzyme B, (**C**), CD107a, (**D**) CD158a, and (**E**) CD38. (**F**) Heatmap of activated cytotoxic NK cell ($CD16^{+}$ $CD56^{dim}$) percent positive protein markers after a 26-hour culture of PBMC with TCM or positive (+) controls evaluated by flow cytometry and percentages of (**G**) Granzyme B, (**H**), CD107a, (**I**) CD158a, and (**J**) CD38. (**K**) Heatmap of unactivated cytokine-secreting NK cell ($CD16^{-}$ $CD56^{++}$) percent positive protein markers after a 26-hour culture of PBMC with TCM or positive (+) controls evaluated by flow cytometry and percentages of (**L**) Granzyme B, (**M**), CD107a, (**N**) CD158a, and (**O**) CD38. (**P**) Heatmap of activated cytokine-secreting NK cell ($CD16^{-}$ $CD56^{++}$) percent positive protein markers after a 26-hour culture of PBMC with TCM or positive (+) controls evaluated by flow cytometry and percentages of (**Q**) Granzyme B, (**R**), CD107a, (**S**) CD158a, and (**T**) CD38 from $CD16^{-}$ $CD56^{++}$ cells within unactivated PBMCs. (**U-V**) Percentage of $CD16^{+}$ (left) and $CD56^{+}$ (right) live cells in (**U**) unactivated and (**V**) activated conditions. Data are shown as mean ± SEM and analyzed by ordinary two-way ANOVA with Dunnett's multiple comparisons test to the 2D or media control (*) or ordinary two-way ANOVA with main effects only and Tukey's multiple comparisons test (c and s significant to CT and ST, respectively). * $p < 0.05$, ** $p < 0.01$, *** $p < 0.001$, **** $p < 0.0001$. n=12 from 4 independent experiments.

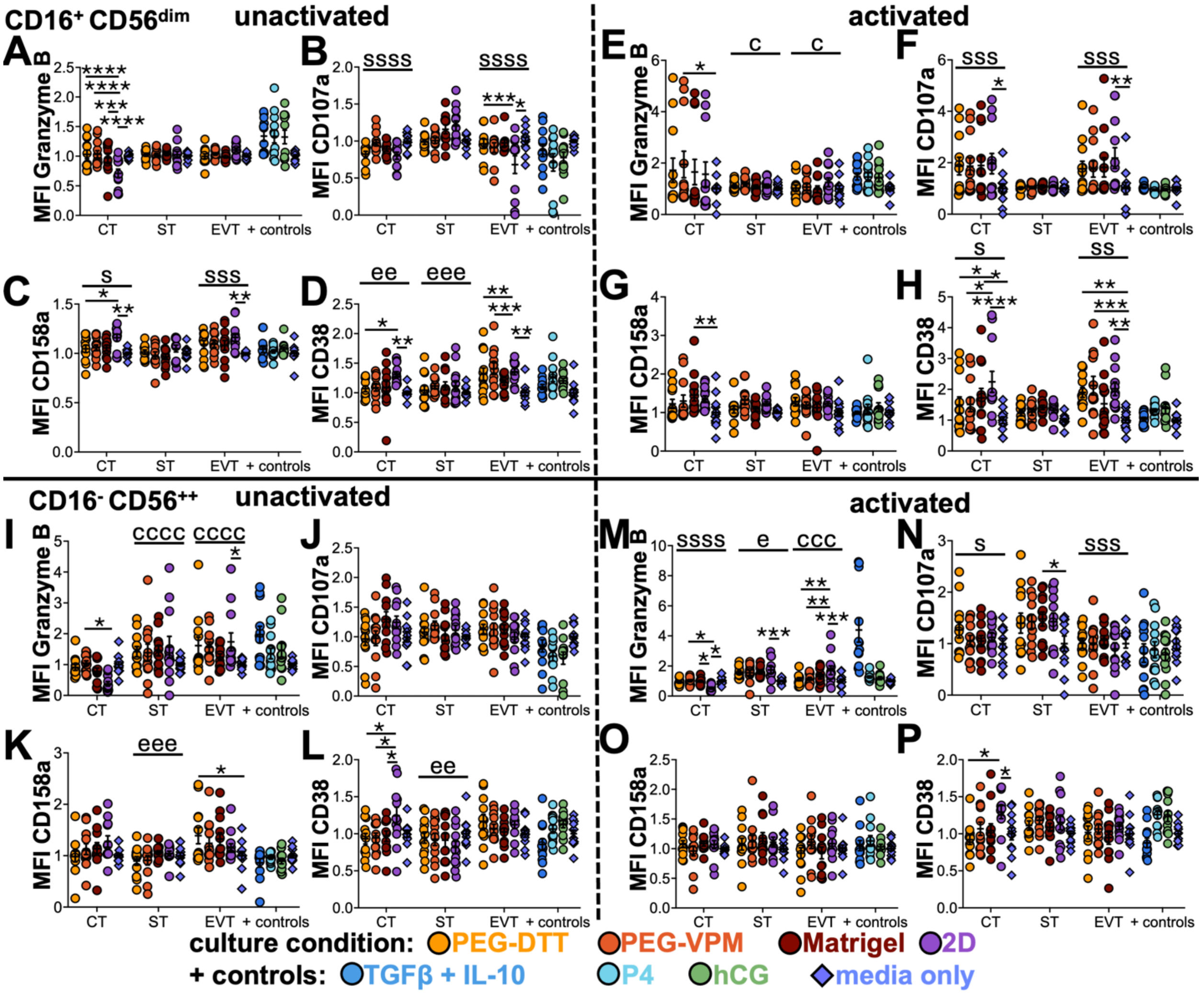


**Fig. S8. CT and EVT TCM significantly modulated NK cell protein MFI within PBMCs**. (**A-D, I-L**) Unactivated and (**E-H, M-P**) activated NK populations within PBMCs cultured with CT, ST,

EVT, and positive controls with protein expression markers assessed via flow cytometry. (**A-H**) $CD16^{+}$ $CD56^{dim}$ and (**I-P**) $CD16^{-}$ $CD56^{++}$ gated populations with markers of (**A, E, I, M**) Granzyme B, (**B, F, J, N**) CD107a, (**C, G, K, O**) CD158a, and (**D, H, L, P**) CD38 normalized MFI to media controls. Data are shown as mean ± SEM and analyzed by ordinary two-way ANOVA with Dunnett's multiple comparisons test to the 2D or media (*) controls or 2D and ordinary two-way ANOVA with main effects only and Tukey's multiple comparisons test (c, s, and e, significant to CT, ST, and EVT, respectively). * $p < 0.05$, ** $p < 0.01$, *** $p < 0.001$, **** $p < 0.0001$. n=12 from 4 independent experiments.

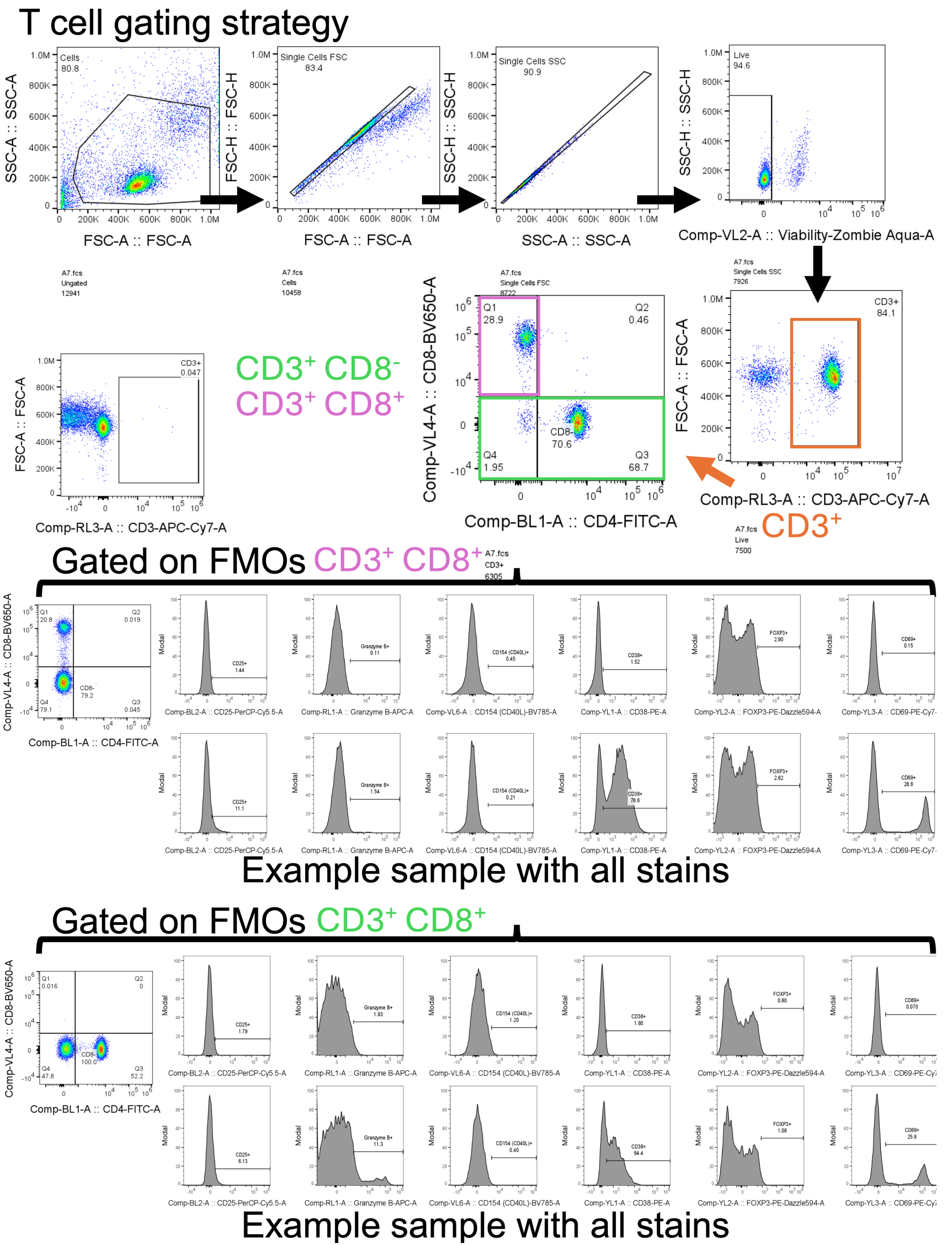


**Fig. S9. T cell gating strategy from PBMC population.** Debris was gated out, singlets, live, and $CD3^+$ events were gated on, then $CD8^+$ and $CD8^-$ cells were assessed for protein marker MFI and percentage of positive cells of protein markers, where fluorescence minus one (FMO) controls were used to gate out cells negative for markers.

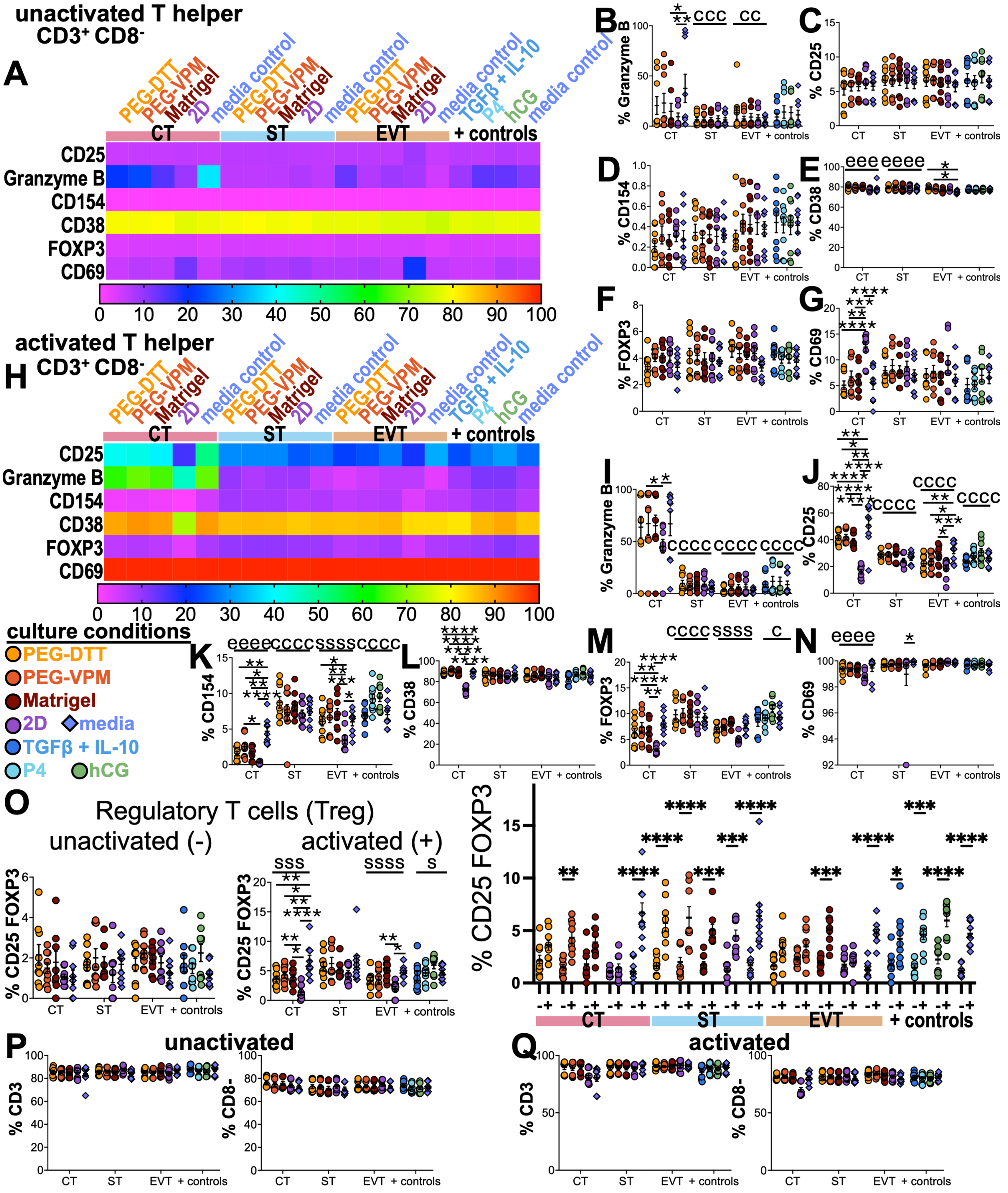


**Fig. S10. CT secretome modulated percent positive activation markers from T helper cells within PBMCs.** (**A, H**) Heatmap of percentage positive protein markers for (**A**) unactivated and (**H**) activated T helper cells ($CD3^{+}$ $CD8^{-}$) after a 48-hour culture of PBMC with TCM or positive (+) controls evaluated by flow cytometry and percentages of (**B, I**) Granzyme B, (**C, J**), CD25, (**D, K**) CD154, (**E, L**) CD38, (**F, M**) FOXP3, and (**G, N**) CD69 positive cells. (**P-Q**) Percentage of $CD3^{+}$ (left) and $CD3^{+}$ $CD8^{-}$ (right) live cells in (**P**) unactivated and (**Q**) activated conditions. Data are shown as mean ± SEM and analyzed by ordinary two-way ANOVA with Dunnett's multiple comparisons test to the media or 2D (*) control or ordinary two-way ANOVA with main effects only and Tukey's multiple comparisons test (c, s, and e, significant to CT, ST, and EVT, respectively). * $p < 0.05$, ** $p < 0.01$, *** $p < 0.001$, **** $p < 0.0001$. n=9 from 3 independent experiments.

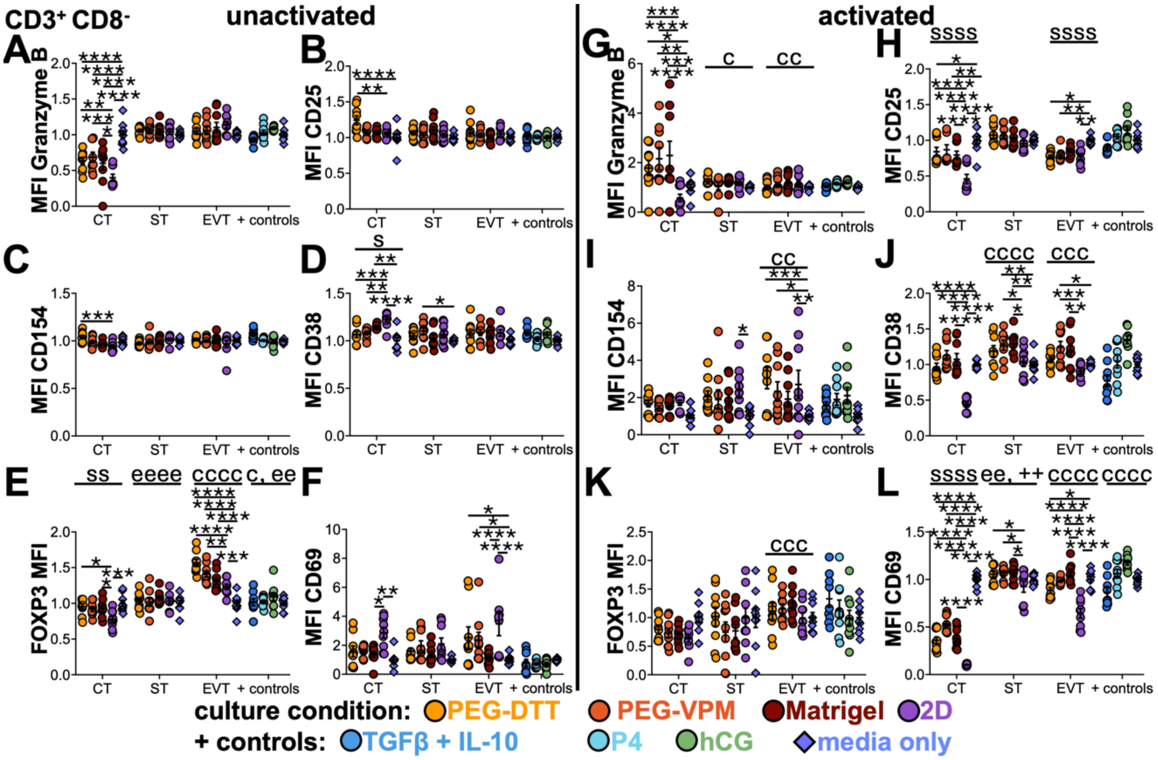


**Fig. S11. CT, ST, and EVT secretome significantly modulated T helper cell protein MFI within PBMCs**. (**A-F**) Unactivated and (**G-L**) activated PBMCs cultured with CT, ST, EVT, and positive controls with protein expression markers assessed via flow cytometry. (**A-H**) $CD3^+$ $CD8^-$ gated populations with markers of (**A, G**) granzyme B, (**B, H**) CD25, (**C, I**) CD154, (**D, J**) CD38, (**E, K**) FOXP3, and (**F, L**) CD69 MFI. MFIs are normalized to media-specific controls. Data are shown as mean ± SEM and analyzed by ordinary two-way ANOVA with Dunnett's multiple comparisons test to the 2D or media (*) control and ordinary two-way ANOVA with main effects only and Tukey's multiple comparisons test (c, s, and e, significant to CT, ST, and EVT, respectively). * $p < 0.05$, ** $p < 0.01$, *** $p < 0.001$, **** $p < 0.0001$. n=9 from 3 independent experiments.

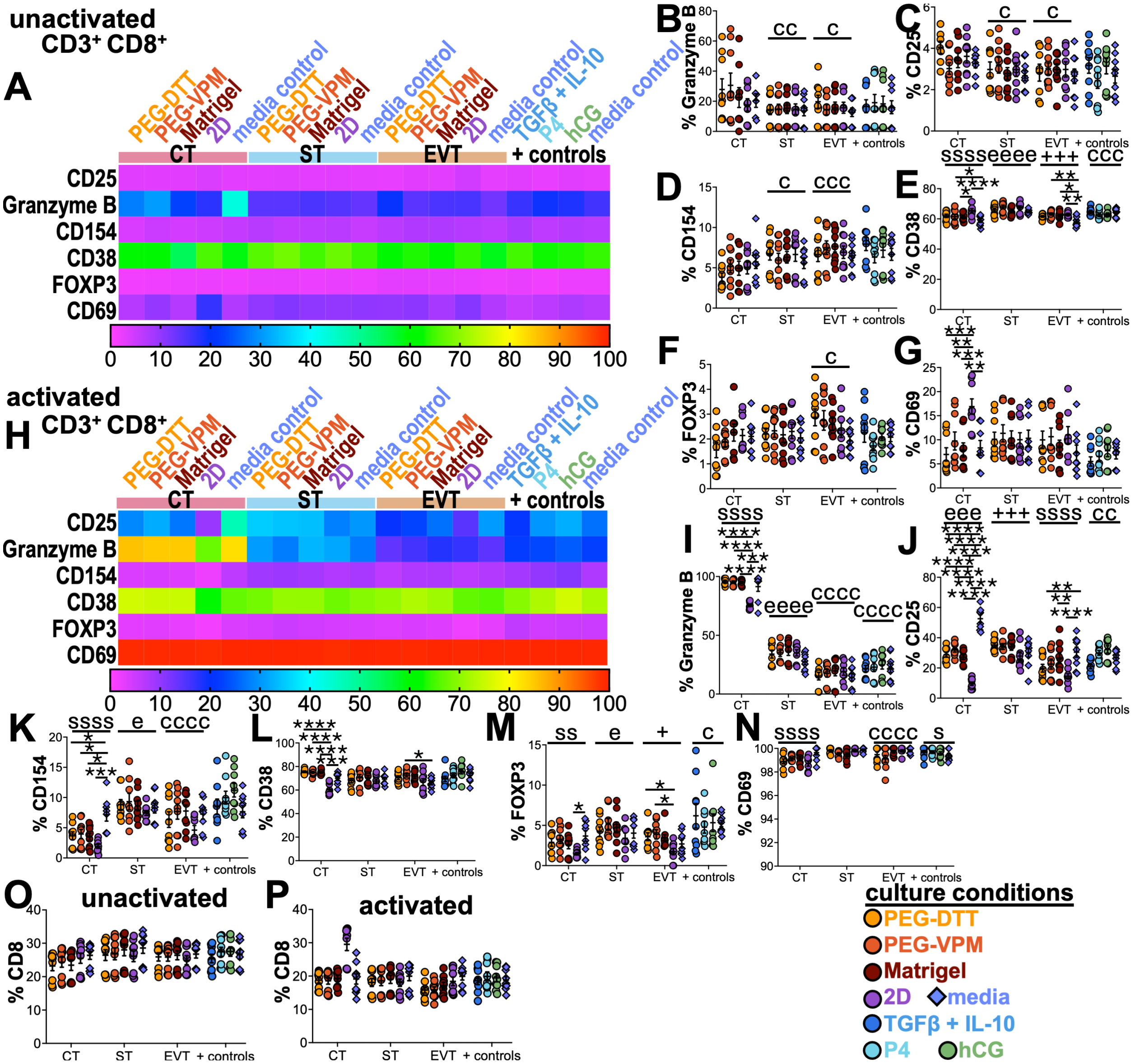


**Fig. S12. CT 2D secretome significantly decreased the percentage of cytotoxic T cells with positive activation markers in activated conditions within PBMCs.** (**A, H**) Heatmap of percentage positive protein markers for (**A**) unactivated and (**H**) activated cytotoxic T cells ($CD3^+$ $CD8^+$) after a 48-hour culture of PBMC with TCM or positive (+) controls evaluated by flow cytometry and percentages of (**B, I**) Granzyme B, (**C, J**), CD25, (**D, K**) CD154, (**E, L**) CD38, (**F, M**) FOXP3, and (**G, N**) CD69. (**O-P**) Percentage of $CD8^+$ live cells in (**O**) unactivated and (**P**) activated conditions. Data are shown as mean ± SEM and analyzed by ordinary two-way ANOVA with Dunnett's multiple comparisons test to the media or 2D (*) control or ordinary two-way ANOVA with main effects only and Tukey's multiple comparisons test (c, s, and e, significant to CT, ST, and EVT, respectively). * $p < 0.05$, ** $p < 0.01$, *** $p < 0.001$, **** $p < 0.0001$. n=9 from 3 independent experiments.

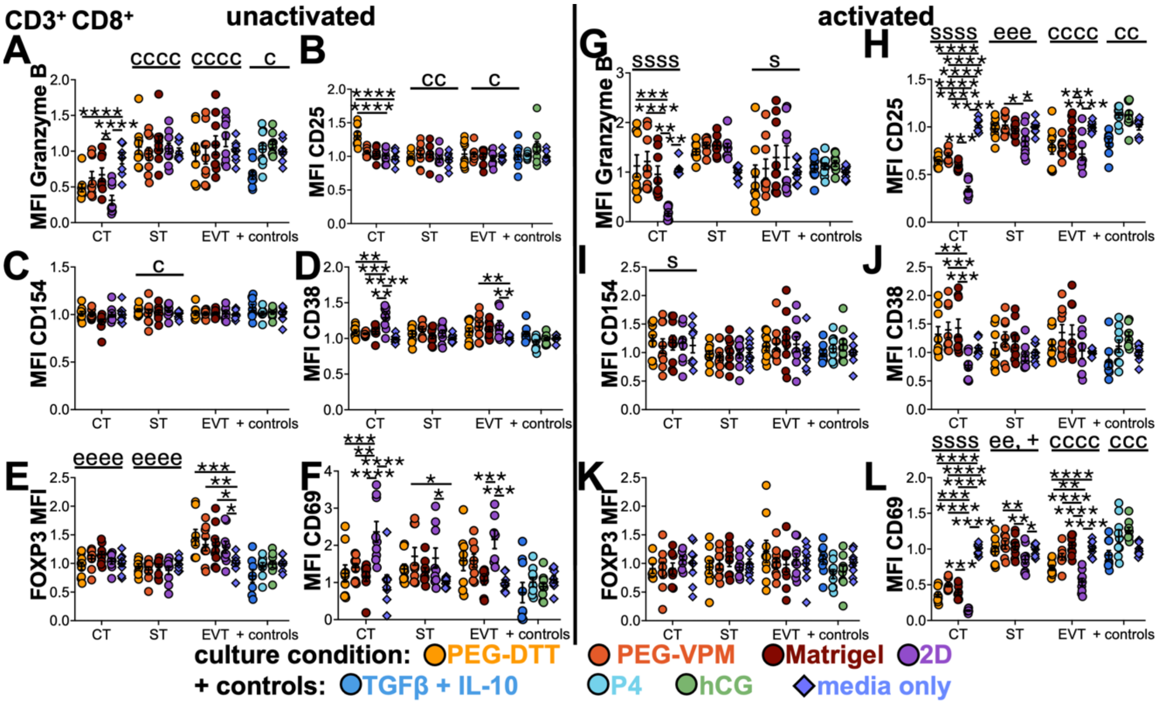


**Fig. S13. CT and EVT secretome differentially modulated cytotoxic T cell protein MFI within PBMCs via unactivated and activated conditions, respectively**. (**A-F**) Unactivated and (**G-L**) activated PBMCs cultured with CT, ST, EVT, and positive controls with protein expression markers assessed via flow cytometry. (**A-H**) CD3$^+$ CD8$^+$ gated populations with markers of (**A, G**) granzyme B, (**B, H**) CD25, (**C, I**) CD154, (**D, J**) CD38, (**E, K**) FOXP3, and (**F, L**) CD69 normalized MFI to respective media controls. Data are shown as mean ± SEM and analyzed by ordinary two-way ANOVA with Dunnett's multiple comparisons test to the media control or 2D (*) and ordinary two-way ANOVA with main effects only and Tukey's multiple comparisons test (c, s, and e, significant to CT, ST, and EVT, respectively). * $p < 0.05$, ** $p < 0.01$, *** $p < 0.001$, **** $p < 0.0001$. n=9 from 3 independent experiments.

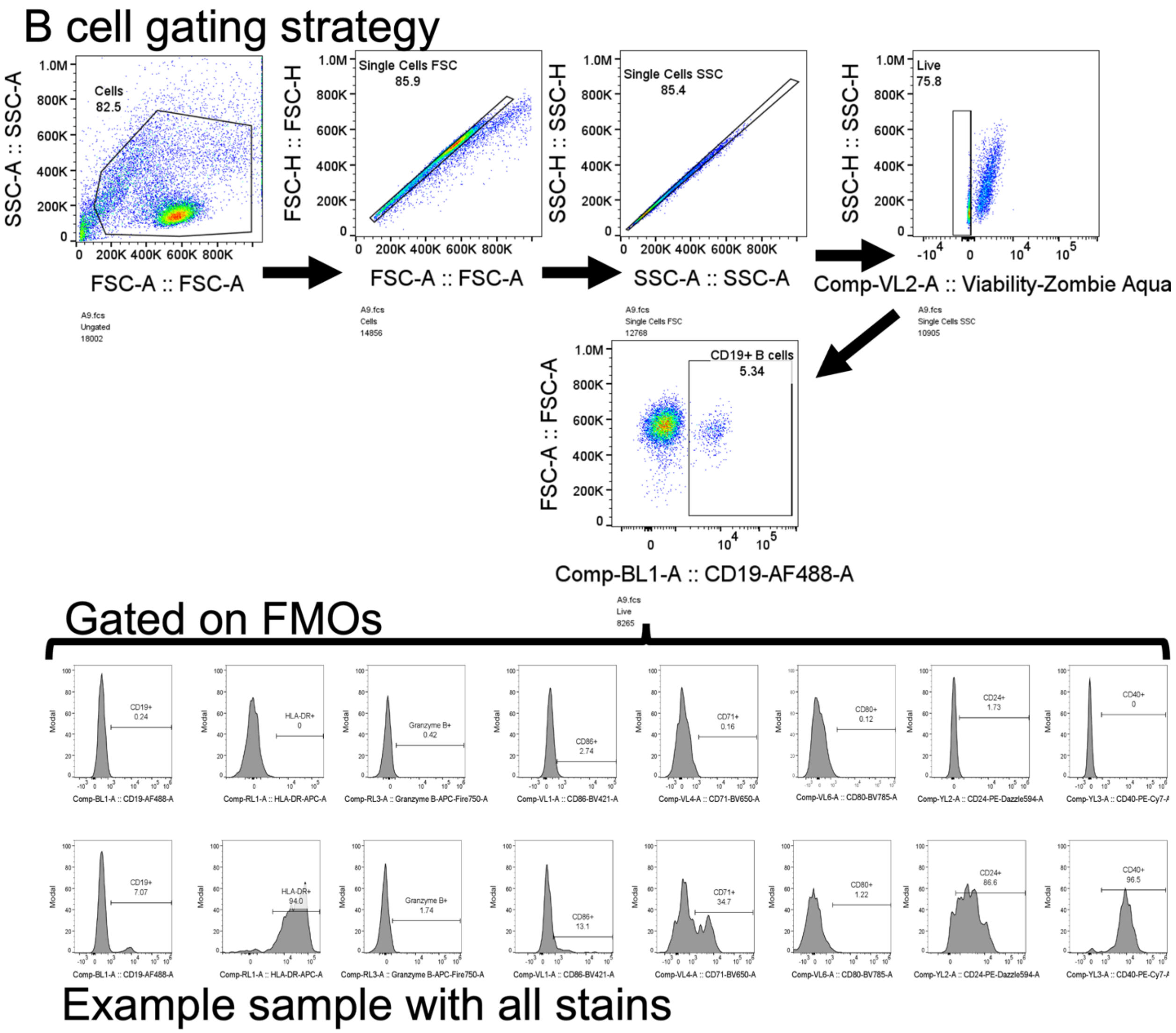


**Fig. S14. B cell gating strategy from PBMC population.** Debris was gated out, singlets and live events were gated on, and then $CD19^{+}$ events were assessed for MFI and percentage of positive cells of protein markers, where fluorescence minus one (FMO) controls were used to gate out cells negative for markers.

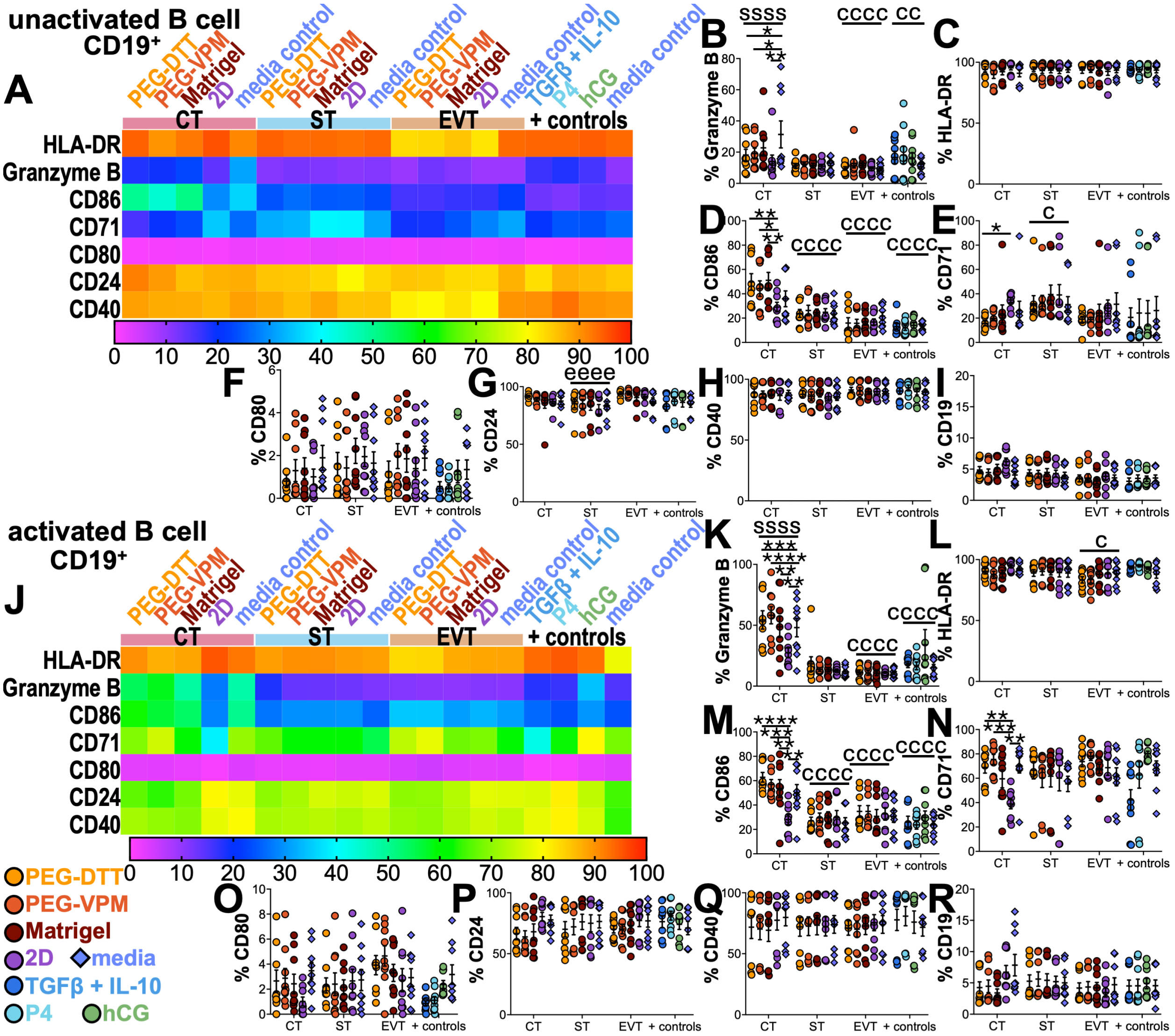


**Fig. S15. CT secretome modulated percentages of B cells positive for phenotypic markers within PBMCs.** (**A, J**) Heatmap of percentage positive protein markers for (**A**) unactivated and (J) activated B (CD19+) cells after a 48-hour culture of PBMC with TCM or positive (+) controls evaluated by flow cytometry and percentages of (**B, K**) Granzyme B, (**C, L**), HLA-DR, (**D, M**) CD86, (**E, N**) CD71, (**F, O**) CD80, (**G, P**) CD24, (**H, Q**) CD40, and (**I, R**) CD19 positive live cells. Data are shown as mean ± SEM and analyzed by ordinary two-way ANOVA with Dunnett's multiple comparisons test to the media or 2D (*) control or ordinary two-way ANOVA with main effects only and Tukey's multiple comparisons test (c, s, and e, significant to CT, ST, and EVT, respectively). * $p < 0.05$, ** $p < 0.01$, *** $p < 0.001$, **** $p < 0.0001$. n=9 from 3 independent experiments.

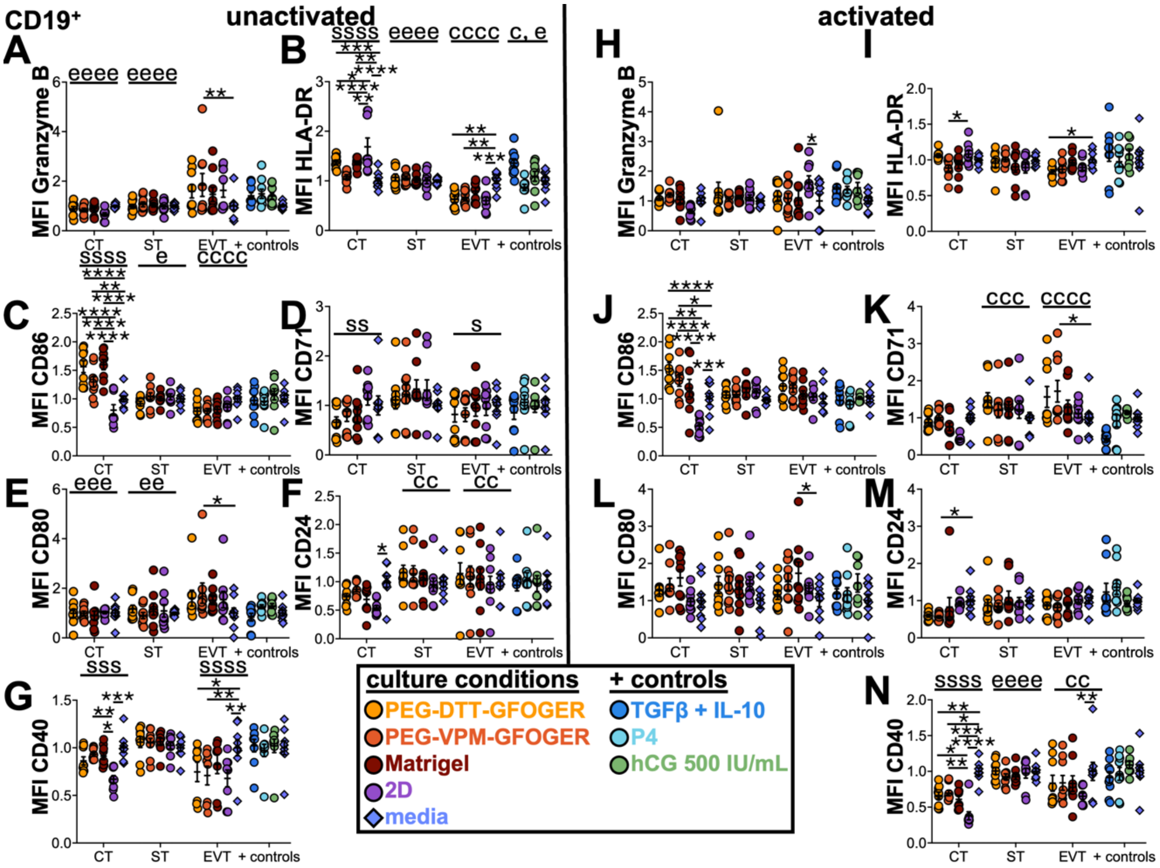


**Fig. S16. CT TCM significantly modulated B cell phenotypic markers within PBMCs**. (**A-G**) Unactivated and (**H-N**) activated PBMCs cultured with CT, ST, EVT, and positive controls with protein expression markers assessed via flow cytometry. CD19+ gated populations with markers of (**A, H**) Granzyme B, (**B, I**) HLA-DR, (**C, J**) CD86, (**D, K**) CD71, (**E, L**) CD80, (**F, M**) CD24, and (**G, N**) CD40 MFI. Data are shown as mean ± SEM and analyzed by ordinary two-way ANOVA with Dunnett's multiple comparisons test to the media control or 2D and ordinary two-way ANOVA with main effects only and Tukey's multiple comparisons test (c, s, and e, significant to CT, ST, and EVT, respectively). * $p < 0.05$, ** $p < 0.01$, *** $p < 0.001$, **** $p < 0.0001$. n=9 from 3 independent experiments.
